# A Comprehensive Immunologic Portrait of Triple-Negative Breast Cancer

**DOI:** 10.1101/209288

**Authors:** Zhixian Liu, Mengyuan Li, Zehang Jiang, Xiaosheng Wang

## Abstract

**Background:** Triple-negative breast cancer (TNBC) is a high-risk malignancy due to its high capacity for invasion and lack of targeted therapy. Immunotherapy continues to demonstrate efficacy in a variety of cancers, and thus may be a promising strategy for TNBC given the limited therapeutic options currently available for TNBC. In this study, we performed a comprehensive portrait of immunologic landscape of TNBC based on 2 large-scale breast cancer genomic data.

**Methods:** We compared expression levels of immune-related genes and gene-sets among TNBC, non-TNBC, and normal tissue, and within TNBCs of different genotypic or phenotypic features. Moreover, we explored the association of immune-related genes or gene-sets expression and survival prognosis in TNBC patients.

**Results:** We found that almost all analyzed immune-related gene-sets had significantly higher expression levels in TNBC than non-TNBC. These highly expressed gene-sets in TNBC included 15 immune cell type and function, human leukocyte antigen (HLA), cancer testis (CT), tumor-infiltrating lymphocytes (TILs), immune cell infiltrate, regulatory T (Treg) cells, immune checkpoint, cytokine and cytokine receptor, metastasis-promoting, pro-inflammatory and parainflammation (PI) gene-sets. Moreover, *TP53-mutated*, TNBC had significantly higher expression levels of the immune checkpoint, Treg, PI, and CT gene-sets, and lower expression levels of the immune cell infiltrate gene-set than *TP53*-wildtype TNBC. Furthermore, we found that elevated expression of most of the immune-related genes in TNBC was associated with the ER-status, while some were associated with both ER-and HER2-status. Elevated expression of the immune-related genes in TNBC was also associated with the high tumor mutation burden (TMB) in TNBC. Finally, elevated expression of the immune-related gene-sets was likely to be associated with better survival prognosis in TNBC.

**Conclusions:** Our findings suggest that TNBC is a breast cancer subtype with particularly strong immunogenicity, and therefore could be propitious to immunotherapeutic options.

## Background

Breast cancer (BC) is the most common cancer in women [1], of which 15-20% are the triplenegative breast cancer (TNBC) subtype. TNBC is clinically negative for expression of the estrogen receptor (ER) and progesterone receptor (PR), and lacks overexpression of the human epidermal growth factor receptor 2 (HER2) [2]. TNBC has a poor prognosis due to its aggressive clinical characteristics and lack of response to hormonal or HER2 receptor-targeted therapy. Thus far, chemotherapy is the only possible therapeutic strategy in the adjuvant or metastatic setting for TNBC [3]. Some potential targeted therapies for TNBC have been investigated such as targeting VEGF, EGFR, mTOR, PARP1, FGFR, AR, NOTCH, HDAC, CDK, PI3K, MET, and TROP2 [4–7]. However, clinical trial efficacies of most TNBC targeted therapies remain unclear.

Recently, cancer immunotherapy has demonstrated high efficacy in treating a variety of cancers including refractory malignancies such as metastatic melanoma and advanced squamous nonsmall cell lung cancer (NSCLC) [8]. Based on the promising results from these other cancers, immunotherapy for TNBC is a viable clinical objective, especially considering the very limited therapeutic options currently available for TNBC. Consequently, several studies have explored the use of immunotherapy against TNBC [9, 10]. For example, Nanda *et al.* provided preliminary evidence demonstrating that pembrolizumab, a highly selective monoclonal IgG4-k antibody against PD1, may be promising in treating advanced TNBC [9]. Emens *et al.* showed that inhibition of PD-L1 by MPDL3280A had encouraging clinical activity in heavily pretreated metastatic TNBC patients [10]. In addition, Hartman *et al.* demonstrated that combined inhibition of IL-6 and IL-8 might be an effective treatment strategy for TNBC [11].

One of the most exciting advances in the field of cancer immunotherapy has been the blockade of immune checkpoint molecules such as cytotoxic T-lymphocyte-associated protein 4 (CTLA4), programmed cell death protein 1 (PD1), and programmed cell death 1 ligand (PD-L1) [12, 13]. The FDA has recently approved immune checkpoint inhibitors such as ipilimumab (anti-CTLA4), nivolumab and pembrolizumab (anti-PD1), and atezolizumab and avelumab (anti-PD-L1) for the treatment of various advanced malignancies such as melanoma, NSCLC, renal cell cancer, Hodgkin's lymphoma, bladder cancer, and head and neck cancer. However, only a subset of patients can benefit from such therapy, with some patients achieving a limited response or completely failing to respond to such therapy [14]. Thus, it is crucial to identify molecular biomarkers for predicting responders to cancer immunotherapy. Some biomarkers have consequently been explored based on genomic or transcriptomic approaches. For example, several studies have revealed the positive correlation of tumor mutation load with clinical response of cancer patients to CTLA4 or PD1 blockade [15–18]. Le *et al.* showed that high mismatch repair (MMR) deficiency correlated with active clinical response to immune checkpoint blockade in cancers [19]. Allen *et al.* demonstrated that tumor mutation load, neoantigen load, and expression of cytolytic markers in the immune microenvironment correlated with clinical response to CTLA4 blockade in metastatic melanoma [18]. These previous explorations of correlating genomic features with cancer immunotherapy response have provided interesting findings. However, genomic biomarkers for precisely predicting responders to cancer immunotherapy are still lacking. This underscores the need for comprehensive and extensive analyses of cancer genomics profiles to discover immunotherapy-responsive biomarkers.

Although BC does not show high responsiveness to immunotherapy as compared to melanoma, lung cancer, renal cancer, lymphoma, bladder cancer, or head and neck cancer, growing evidence suggests the existence of variable immunogenic activity in BC subtypes [20, 21]. Several studies have identified immunogenic subtypes of BC or TNBC, suggesting that immunogenic heterogeneity may correlate with phenotypic heterogeneity of BC [20–22]. In a recent study [23], Safonov *et al.* analyzed the gene expression, DNA copy number, somatic and germline mutation data of BC from The Cancer Genome Atlas (TCGA), and found that TNBC and HER2+ BC had high immune gene expression and lower clonal heterogeneity as compared to other BC subtypes. Another recent study found a correlation between the expression of immunologic signatures and clinical outcomes in TNBC, and demonstrated that elevated expression of *HLA-C*, *HLA-F*, *HLA-G*, and *TIGIT* were associated with improved relapse-free survival and overall survival (OS) [24].

However, these previous studies only analyzed 1 or several aspects of immune function in TNBC [20–24]. To fill the gaps in knowledge of immunologic landscape of TNBC, we performed a comprehensive and exhaustive analysis of immunogenic signatures in TNBC based on 2 large-scale BC genomics datasets: The Cancer Genome Atlas (TCGA) and METABRIC BC [25-27]. We compared expression of immune-related genes and gene-sets among TNBC, non-TNBC, and normal tissue, and within TNBCs of different genotypes or phenotypes. In addition, we evaluated the degree of immune cell infiltration in different BC subtypes by ESTIMATE [28] and CIBERSORT [29]. Our study aimed to address the following questions, including: Is the immunogenic activity of TNBC different from other BC subtypes? What molecular cues are associated with the differences in the immunogenic activity between TNBC and other BC subtypes? Is tumor mutation load associated with the immunogenic activity of TNBC? Are there any immune-related genes or gene-sets whose expression is associated with clinical outcomes in TNBC?

## Results

### TNBC has higher expression levels of immune cell types and functional marker genes than non-TNBC and normal tissue

We analyzed 15 immune cell types and functional gene-sets associated with B cells, CD4+ regulatory T cells, CD8+ T cells, macrophages, neutrophils, natural killer (NK) cells, plasmacytoid dendritic cells (pDCs), major histocompatibility complex (MHC) class I, APC costimulation, T cell co-stimulation, APC co-inhibition, T cell co-inhibition, Type I IFN response, Type II IFN response, and cytolytic activity, respectively [30]. We found significant differential expression in a substantial number of genes in these 15 gene-sets between TNBC and non-TNBC, and the expression differences were almost commonly identified in both TCGA and METABRIC datasets with identical expression change direction (Figure 1A; Supplementary Table S1). For example, all 10 B cell marker genes (*CD79B, BTLA, FCRL3, BANK1, CD79A, BLK, RALGPS2, FCRL1, HVCN1*, and *BACH2*) were differentially expressed between TNBC and non-TNBC in TCGA, and 9 were differentially expressed between TNBC and non-TNBC in METABRIC, except *FCRL1*, which was not included in the METABRIC gene list. Among the 9 differentially expressed genes identified in both datasets, 8 were more highly expressed in TNBC than in non-TNBC. In the 7 CD4+ regulatory T cell marker genes, *C15orf53*, *CTLA4*, and *IL32* were more highly expressed in TNBC than in non-TNBC in both datasets, and *FOXP3* and *GPR15* were more highly expressed in TNBC than in non-TNBC in TCGA. The CD8+ T cell marker gene *CD8A* was more highly expressed in TNBC than in non-TNBC in both datasets. Both NK cell marker genes, *KLRF1* and *KLRC1*, were more highly expressed in TNBC than in non-TNBC in both datasets. Both cytolytic activity marker genes, *GZMA* and *PRF1*, were more highly expressed in TNBC than in non-TNBC in both datasets. Furthermore, the majority of macrophages, MHC class I, APC co-stimulation, T cell co-stimulation, APC co-inhibition, and T cell co-inhibition marker genes were more highly expressed in TNBC than in non-TNBC in both datasets. In the TCGA dataset with normal controls, a large number of immune cell types and functional genes also had significantly higher expression levels in TNBC than in normal tissue.

**Figure 1.**
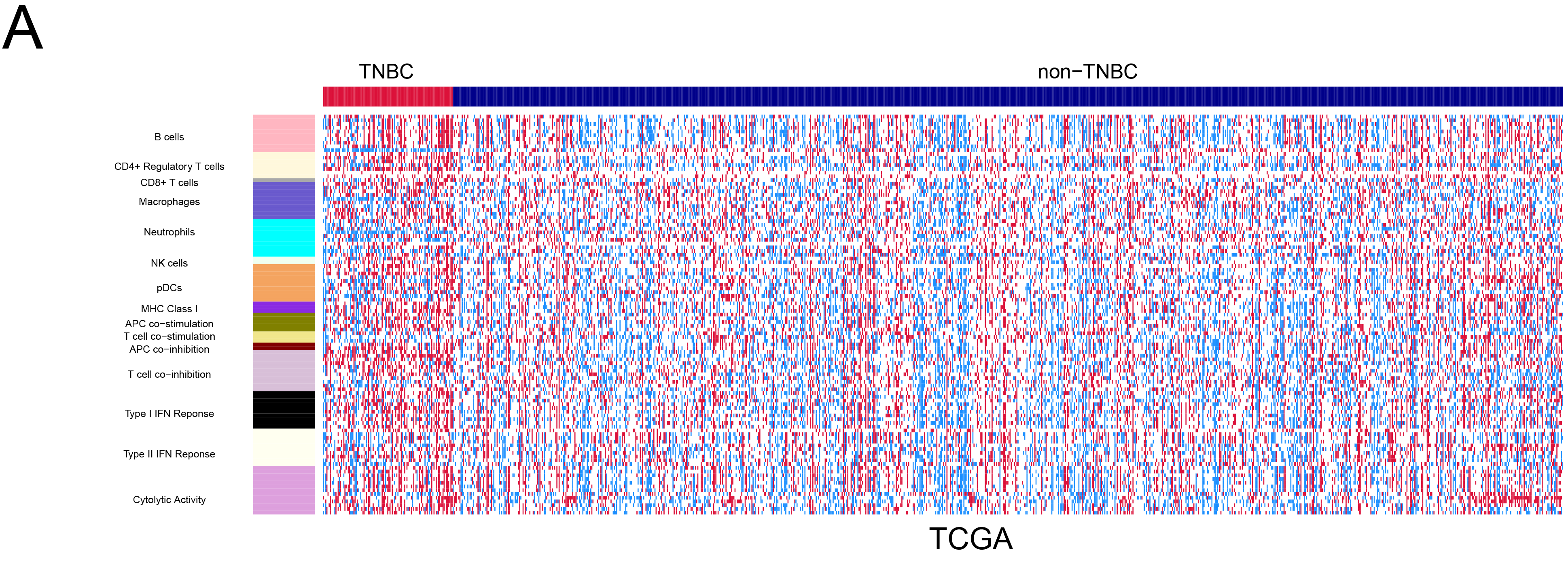
Comparison of expression levels of immune cell types, functional markers, and HLA genes between TNBC and non-TNBC. **A.** Heat-map for expression levels of immune cell types and function genes in TNBC and non-TNBC. **B.** Comparison of expression levels of the HLA gene-set between TNBC and non-TNBC. Red color indicates higher gene expression levels, and blue color indicates lower gene expression levels.

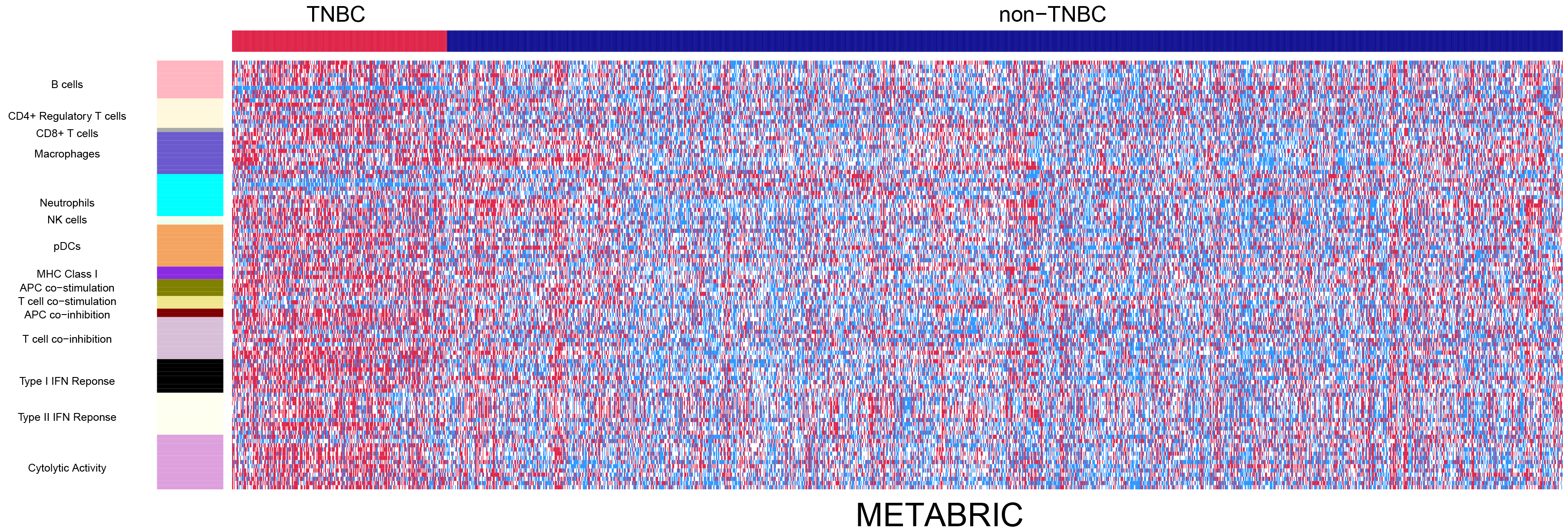

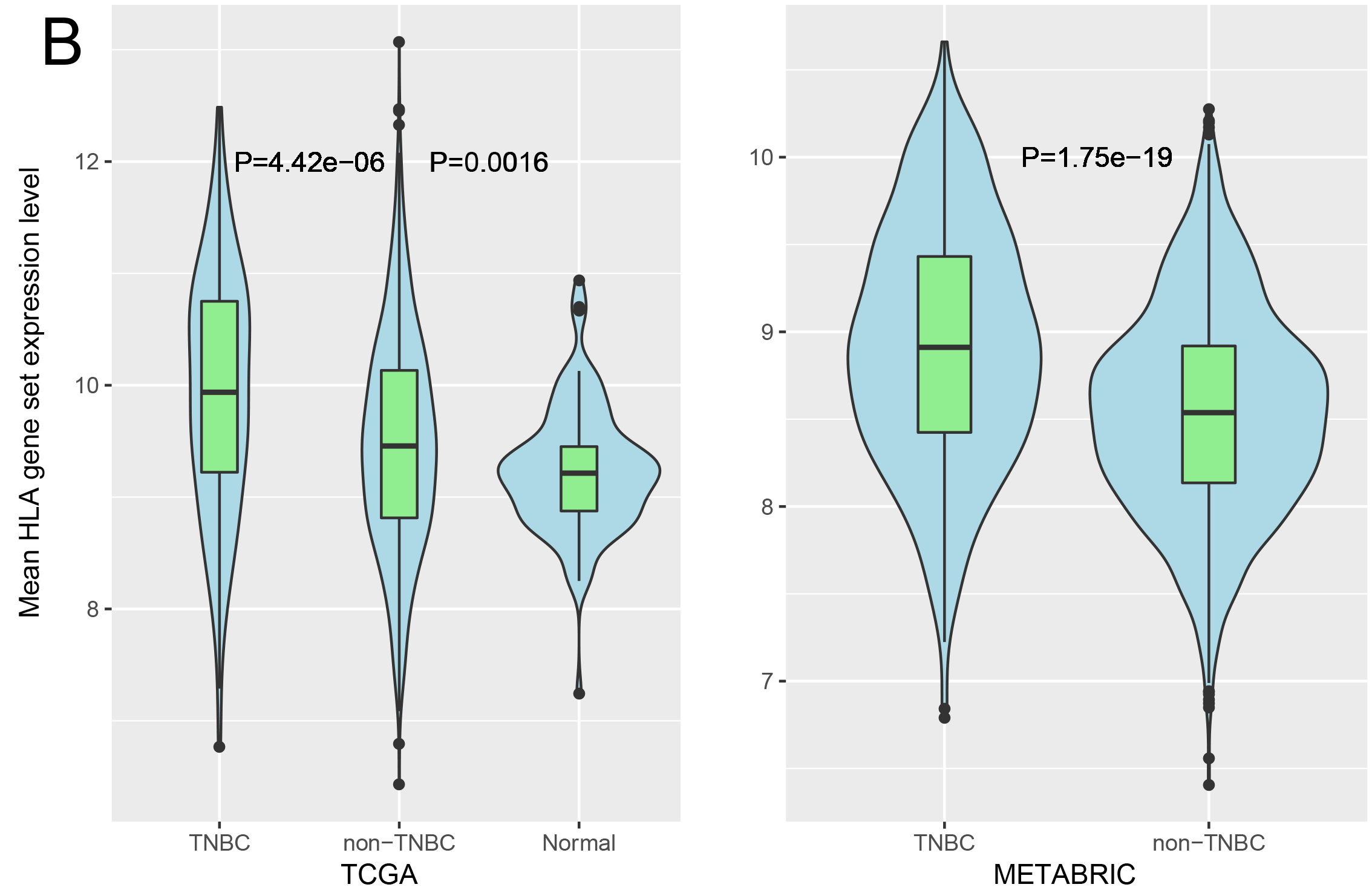

We quantified the activity of an immune cell type or function as the mean expression levels of the respective genes. Interestingly, all 15 immune cell types and functional markers showed higher activities in TNBC than in non-TNBC in METABRIC, and 12 in TCGA (Wilcox rank-sum test, FDR<0.05) (Supplementary Figure S1A, Table S1). Moreover, 10 immune cell types and functional markers had higher activities in TNBC than in normal tissue (Supplementary Figure S1A, Table S1). These results suggest that TNBC likely had elevated immunogenic activity compared to non-TNBC and normal tissue, a finding that is in line with previous studies [22, 23].

### TNBC shows significant differences in HLA genotypes and phenotypes compared to non-TNBC

HLA genes encode MHC proteins, which are responsible for the regulation of the immune system. We compared HLA genotypes (DNA somatic mutations) and phenotypes (mRNA gene expression) between TNBC and non-TNBC. TCGA data showed that TNBC had higher somatic mutation rates of HLA genes than non-TNBC (Fisher's exact test, P=0.04, OR=1.78), while METABRIC had no somatic mutation data available for HLA genes. Strikingly, most HLA genes showed markedly higher expression levels in TNBC than in non-TNBC in both datasets (Supplementary Table S2). Meanwhile, most HLA genes showed significantly higher expression levels in TNBC than in normal tissue. The expression levels of the HLA gene-set were significantly higher in TNBC than in non-TNBC in both datasets (Wilcox rank-sum test, P=4.42*10^−6^, 1.75*10^−19^ for TCGA and METABRIC, respectively) (Figure 1B). Moreover, both TNBC and non-TNBC had significantly higher expression levels of the HLA gene-set than normal tissue (Wilcox rank-sum test, P=5.8*10^−9^, 1.62*10^−3^ for TNBC and non-TNBC, respectively) (Figure 1B).

Gene mutations may yield neoepitopes that can be recognized by immune cells [31]. We compared total mutation counts between TNBC and non-TNBC in TCGA, and found that TNBC had higher mutation counts than non-TNBC (Wilcox rank-sum test, P=4.39*10^−11^). Moreover, TNBC had significantly higher tumor mutation burden (TMB) than non-TNBC (Wilcox rank-sum test, P=2.2*10^−11^). Rooney *et al.* [30] predicted that mutations introduced novel peptides loading in imputed HLA alleles in TCGA samples. We found that TNBC had more gene mutations yielding predicted HLA-binding peptides than non-TNBC (Wilcox rank-sum test, P=2.01*10^−7^).

Altogether, these results suggest that TNBC has more somatic mutations in HLA genes, higher expression levels of HLA genes, and more gene mutations possibly yielding HLA-binding peptides than non-TNBC, which is indicative of stronger immunogenic activity in TNBC relative to non-TNBC.

### TNBC has higher expression levels of many cancer-testis antigen genes than non-TNBC

Cancer-testis (CT) antigens are immunogenic proteins that are normally expressed only in the human germ line; however, the CT antigens are aberrantly activated and expressed in various cancer types, and therefore are potential targets for therapeutic cancer vaccines [32]. We obtained 233 CT genes from the database *CTdatabase* [33], and examined their expression in both datasets. We found that 63 CT genes were more highly expressed in TNBC than in non-TNBC in both datasets versus 20 CT genes that were more highly expressed in non-TNBC than in TNBC (Fisher's exact test, P=2.21*10^−7^, OR=3.94) (Supplementary Figure S1B, Table S3). Many genes which encode important CT antigens and are potentially useful for developing cancer vaccines were more highly expressed in TNBC than in non-TNBC, including *MAGEA* (*MAGEA-2, 3, 4, 5, 6, 9B, 10, 12*), *NY-ESO-1*, and *PRAME* (Supplementary Figure S1C). The expression levels of the CT gene-set were significantly higher in TNBC than in non-TNBC in both datasets (Wilcox rank-sum test, P=6.02*10^−28^, 1. 14* 10^−35^ for TCGA and METABRIC, respectively). Moreover, both TNBC and non-TNBC had significantly higher expression levels of the CT gene-set than normal tissue (Wilcox rank-sum test, P=7.28*10^−29^, 3.72*10^−7^ for TNBC and non-TNBC, respectively) (Supplementary Figure S1D). The expression levels of the CT gene-set were higher in high-grade TNBC than in low-grade TNBC (Wilcox rank-sum test, P=4.02*10”), indicating that many CT genes have increased expression levels with cancer progression. Interestingly, *TP53*-mutated TNBC had significantly higher expression levels of the CT gene-set than *TP53*-wildtype TNBC in both datasets (Wilcox rank-sum test, P=0.007, 3.42*10^−8^ for TCGA and METABRIC, respectively). These results indicated that p53 might repress the expression of many CT genes, and the loss of repressive function by wildtype p53 may result in the elevated expression of these genes. This finding is consistent with a previous study showing that p53 regulated CT genes [34].

### TNBC has higher degree of immune cell infiltration than non-TNBC

Tumor-infiltrating lymphocytes (TILs) migrate from the bloodstream into the tumor microenvironment (TME). TILs have been associated with cancer prognosis and cancer immunotherapy [35, 36]. We compared expression levels of 122 TILs gene signatures [37] between TNBC and non-TNBC. Strikingly, 113 (93%) TILs genes were more highly expressed in TNBC in at least 1 dataset (91 in both datasets), and only a single gene *GLYR1* was more highly expressed in non-TNBC in both datasets (Figure 2A; Supplementary Table S4). The expression levels of the TILs gene-set were significantly higher in TNBC than in non-TNBC in both datasets (Wilcox rank-sum test, P=2.62*10^−6^, 6.57*10^−29^ for TCGA and METABRIC, respectively). The expression levels of the TILs gene-set were also significantly higher in TNBC than in normal tissue (Wilcox rank-sum test, P=3.43*10^−4^), while showed no significant differences between non-TNBC and normal tissue (Wilcox rank-sum test, P=0.13) (Supplementary Figure S2A).

**Figure 2.**
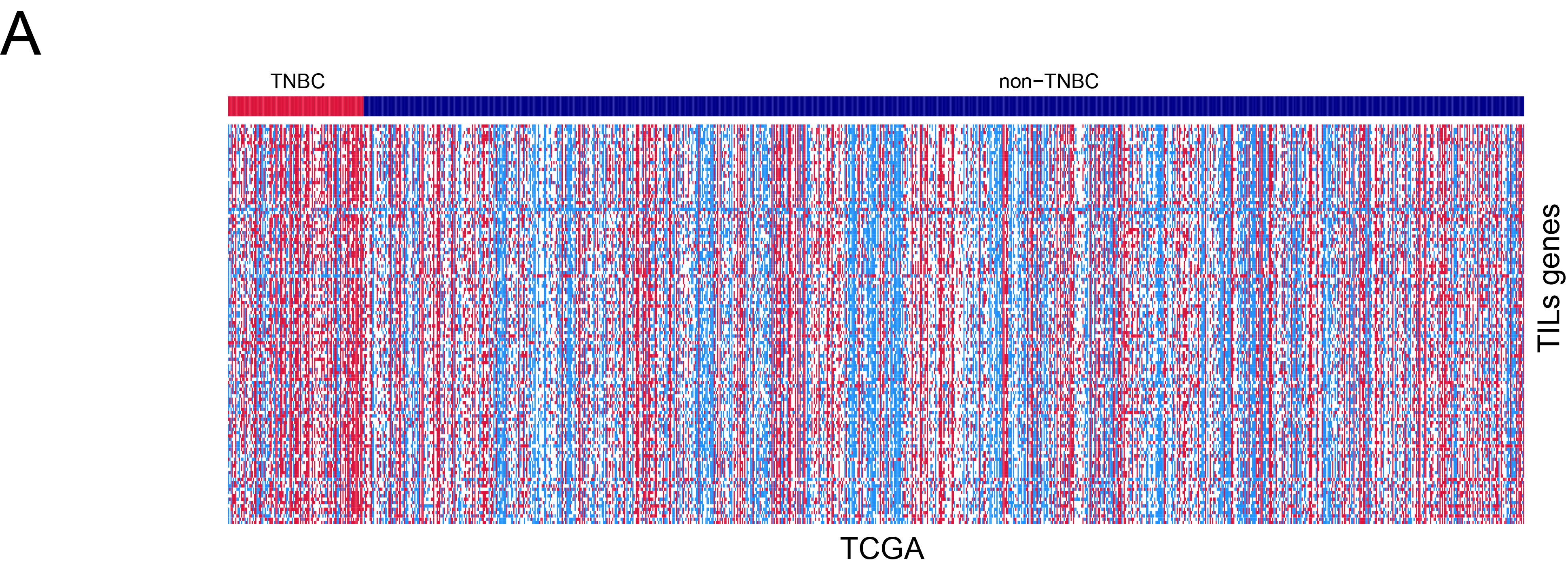
Comparison of expression levels of TILs, immune cell infiltrate, Treg, and immune checkpoint genes between TNBC and non-TNBC. **A.** Heat-map for expression levels of TILs genes in TNBC and non-TNBC. TILs: tumor-infiltrating lymphocytes. **B.** Comparison of expression levels of immune cell subpopulation genes between TNBC and non-TNBC in METABRIC. *: P < 0.05; **: P < 0.01; ***: P < 0.001, and it applies to all the following box charts. **C.** Heat-map for expression levels of Treg and immune checkpoint genes in TNBC and non-TNBC. **D.** Comparison of expression levels of important immune checkpoint genes between TNBC and non-TNBC in METABRIC. Red color indicates higher gene expression levels, and blue color indicates lower gene expression levels.

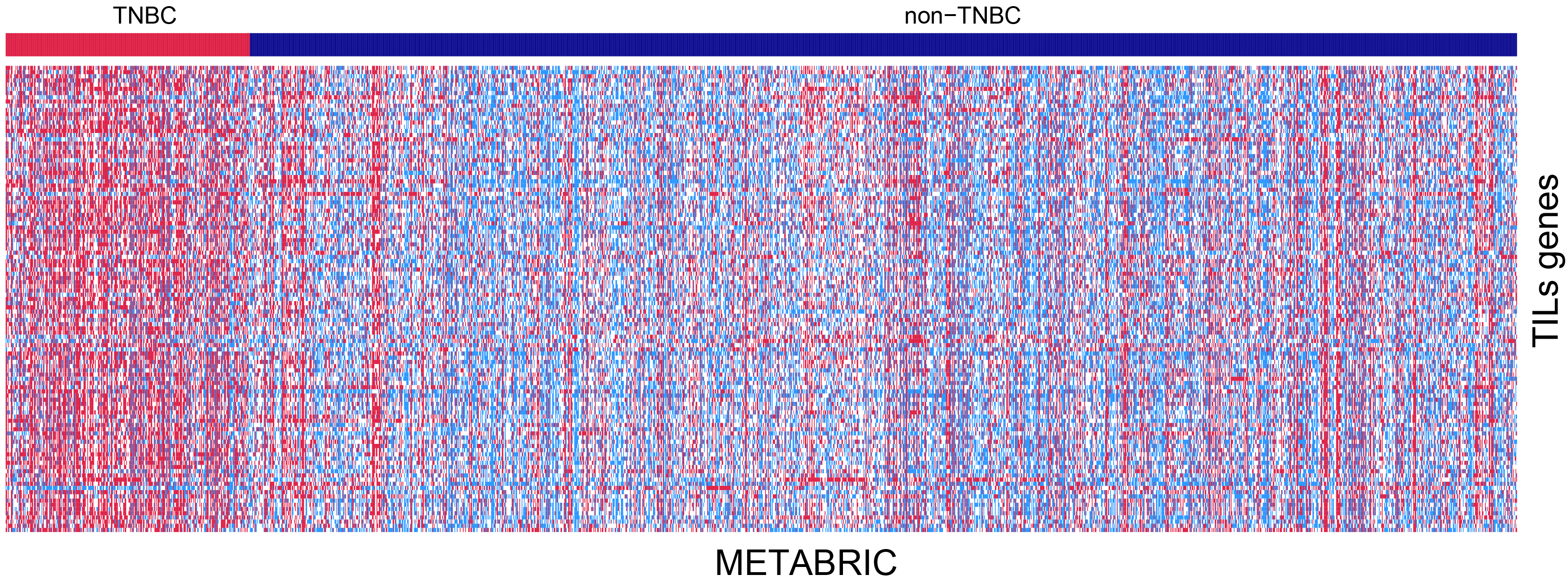

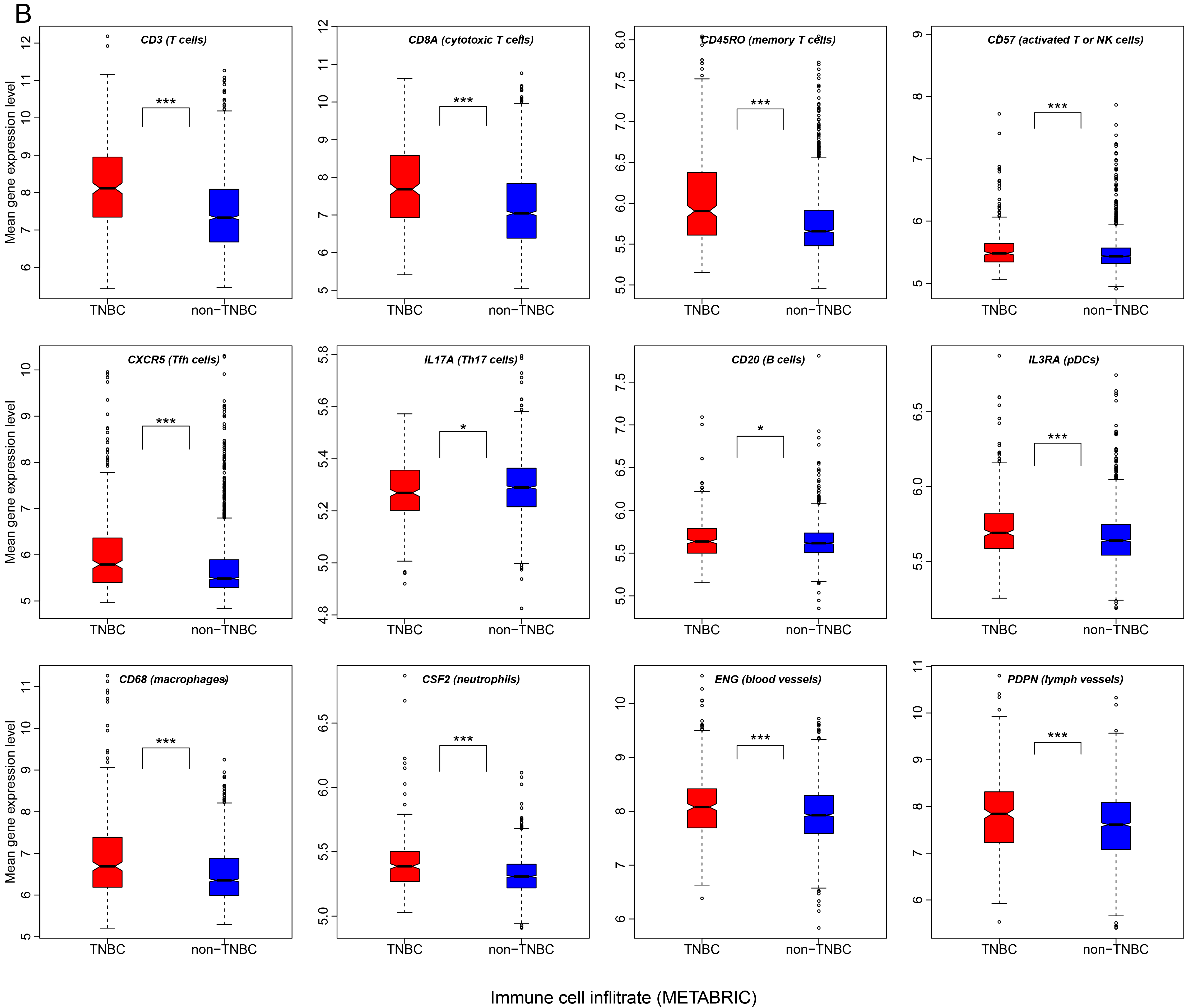

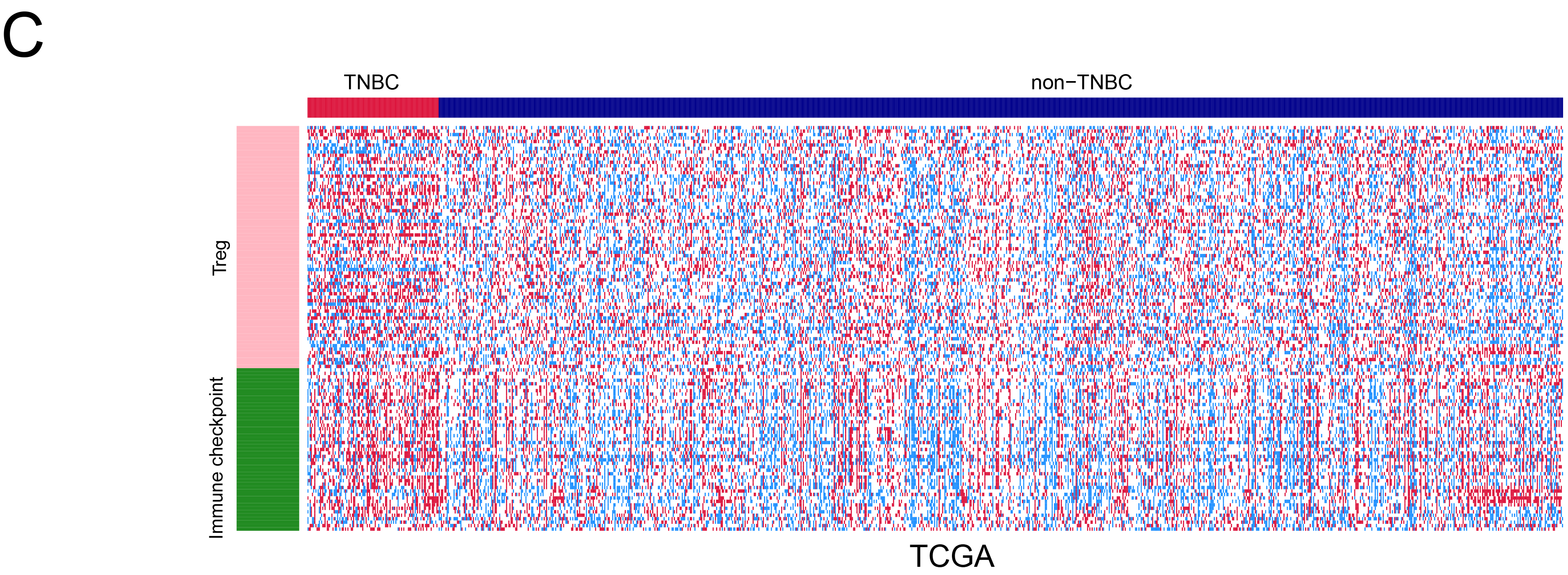

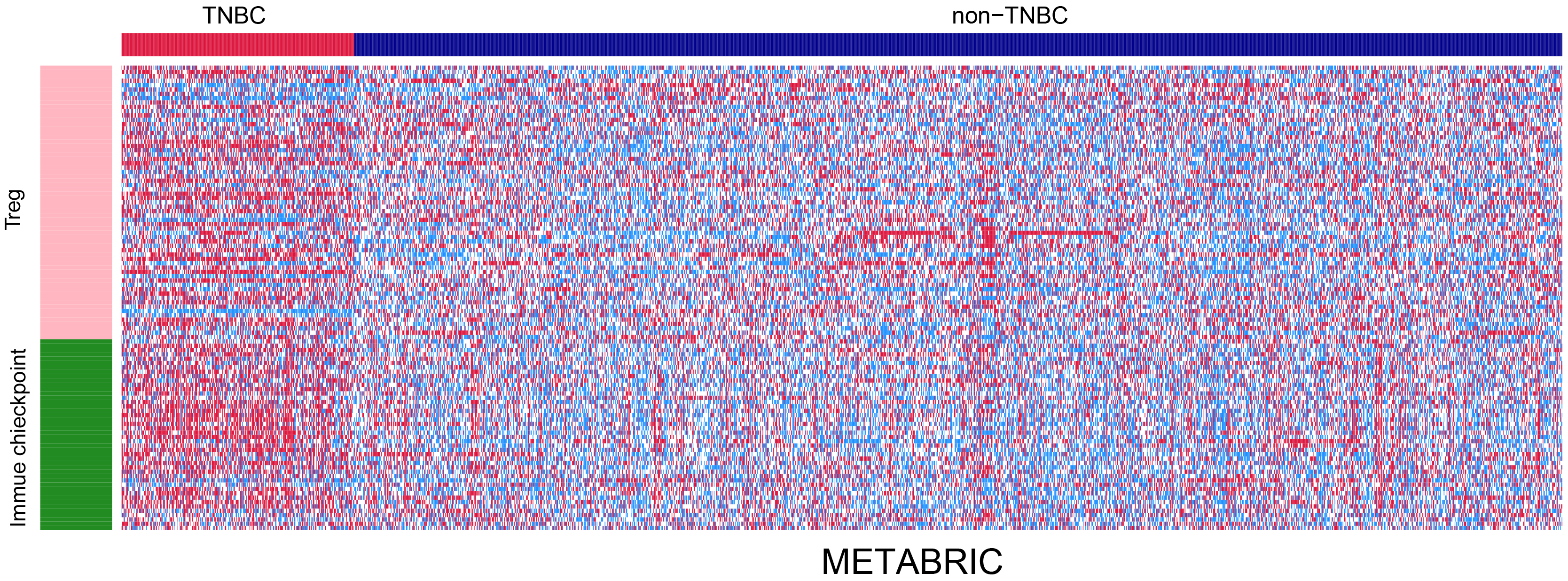

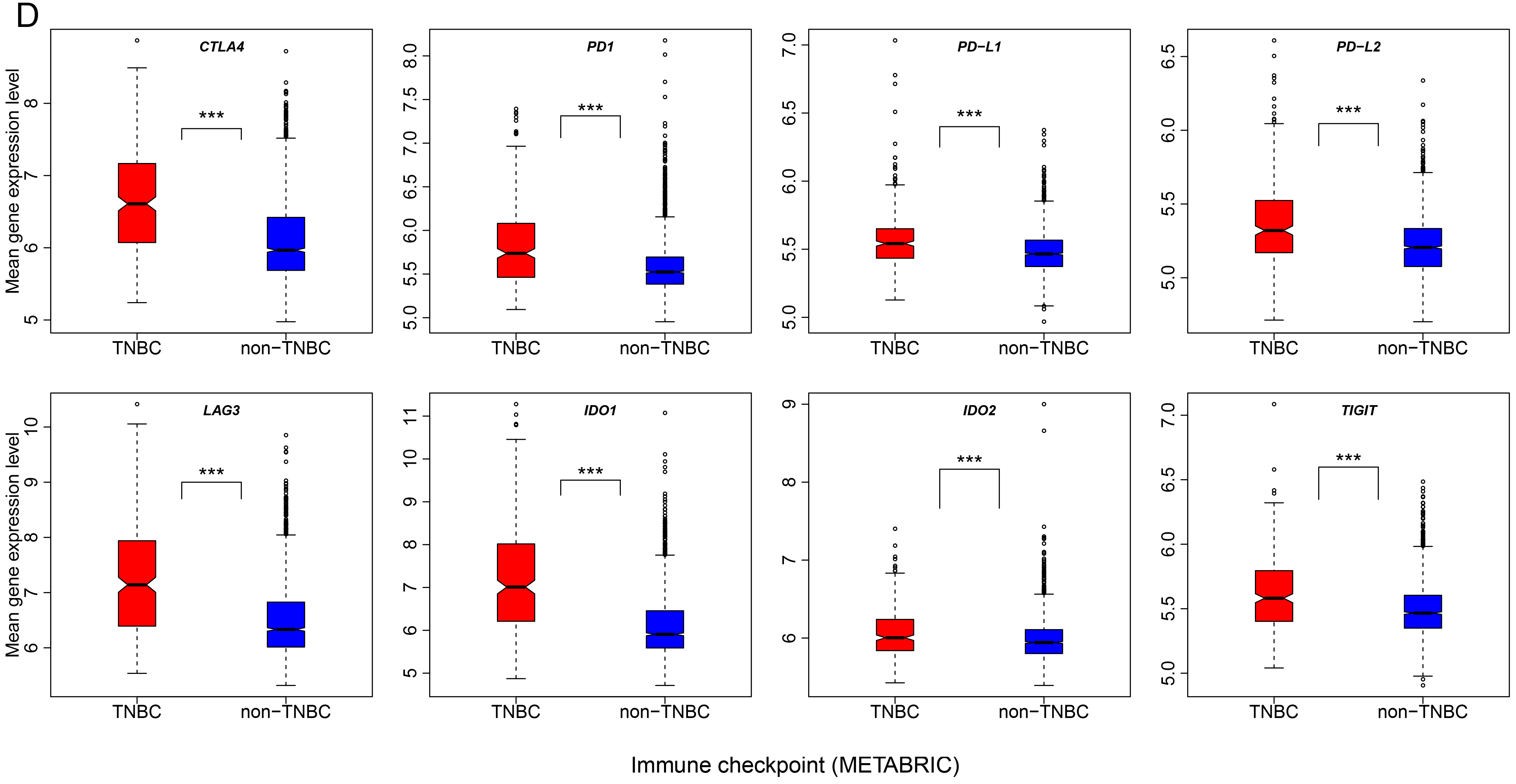

In addition, we compared the immune infiltrate densities of different immune cell subpopulations among TNBC, non-TNBC, and normal tissue. These immune cell subpopulations included T cells (quantified with marker *CD3* gene expression levels), cytotoxic T cells (*CD8*), memory T cells (*CD45RO*), Tregs (*FOXP3*), activated T or NK cells (*CD57*), Tfh cells (*CXCR5*), Th17 cells (*IL-17*), B cells (*CD20*), iDCs (*CD1A*), pDCs (*IL3RA*), macrophages (*CD68*), mast cells (*Tryptase*, *TPSB2*), neutrophils (*CSF2*), blood vessels (*ENG*), and lymph vessels (*PDPN*) [38].

Strikingly, 13 of the 15 immune cell subpopulations marker genes had significantly higher expression levels in TNBC than in non-TNBC in a single or both datasets (Figure 2B; Supplementary Figure S2B, Table S5). This suggests that a higher degree of infiltration occurs in TNBC than in non-TNBC. Interestingly, although *CD57* (activated T or NK cells marker) and *IL3RA* (pDCs marker) were more highly expressed in TNBC than in non-TNBC, both genes had significantly lower expression levels in TNBC and non-TNBC compared to normal tissue. The decreased subpopulations of activated T cells, NK cells, and pDCs in BC suggest the possibility of an immune escape mechanism in BC.

The expression levels of the immune cell infiltrate (subpopulations) marker gene-set were significantly higher in TNBC than in non-TNBC in both datasets (Wilcox rank-sum test, P=5.12*10^−6^, 1.23* 10^−28^ for TCGA and METABRIC, respectively). Interestingly, the expression levels of this gene-set were significantly higher in TNBC than in normal tissue (Wilcox rank-sum test, P=0.008), while significantly lower in non-TNBC than in normal tissue (Wilcox rank-sum test, P=0.006), again demonstrating that TNBC has higher immune cell infiltration levels than non-TNBC. However, no significant differences between low-grade and high-grade TNBC was observed in the expression levels of this gene-set (Wilcox rank-sum test, P=0.18). This data suggests that immune cell infiltrate densities likely do not increase with TNBC progression. Another interesting finding was that *TP53*-mutated TNBC had significantly lower expression levels of this gene-set than *TP53*-wildtype TNBC in METABRIC (Wilcox rank-sum test, P=0.02). This suggests that *TP53* mutations likely influence the levels of immune cell infiltration in TNBC. This finding is in line with the hypothesis that *TP53* mutations may lead to attenuation of immune responses [39].

### TNBC has higher expression levels of immunosuppressive genes than non-TNBC

Regulatory T (Treg) cells are crucial for the maintenance of immunosuppressive activity in cancer [40], and are highly expressed in TNBC [41]. We examined expression levels of 70 tumor-infiltrating Treg (Treg) gene signatures [42] in TNBC. Among the 70 genes, 45 were highly expressed in TNBC as compared to non-TNBC in at least 1 dataset (33 in both datasets) compared to 19 that were more highly expressed in non-TNBC than in TNBC in at least 1 dataset (17 in both datasets) (Fisher's exact test, P=1.8*10^−5^, OR=4.77) (Figure 2C; Supplementary Table S6). The expression levels of the Treg gene-set were significantly higher in TNBC than in non-TNBC in both datasets (Wilcox rank-sum test, P=2.76*10^−10^, 1.84*10^−52^ for TCGA and METABRIC, respectively) (Supplementary Figure S3A). Interestingly, the expression levels of the Treg gene-set were significantly higher in TNBC than in normal tissue (Wilcox rank-sum test, P=6.71*10^−8^), while no significant differences were observed between non-TNBC and normal tissue (Wilcox rank-sum test, P=0.45). Remarkably, *TP53*-mutated TNBC had significantly higher expression levels of the Treg gene-set than *TP53*-wildtype TNBC in TCGA (Wilcox rank-sum test, P=0.01), suggesting that *TP53* mutations may promote Treg infiltration in TNBC.

Immune checkpoints play an important role in tumor immunosuppression [12, 43]. In the 47 immune checkpoint genes provided by De Simone *et al* [42], our study found that 41 (87%) were more highly expressed in TNBC than in non-TNBC in at least 1 dataset (35 in both datasets) versus 4 (9%) that were more highly expressed in non-TNBC than in TNBC in at least 1 dataset (3 in both datasets) (Fisher's exact test, P=2.6*10^−15^, OR=67.23) (Figure 2C; Supplementary Table S7). Moreover, 27 immune checkpoint genes were more highly expressed in TNBC than in normal tissue compared to 9 more highly expressed in normal tissue than in TNBC (Fisher's exact test, P=2.5*10^−4^, OR=5.58). Interestingly, a number of immune checkpoint genes that have been established or considered promising targets for cancer immunotherapy were upregulated in TNBC compared to non-TNBC, and included *CTLA4, PD1, PD-L1, PD-L2, LAG3, IDO1/2*, and *TIGIT.* Of these, *CTLA4*, *PD1*, *LAG3*, *IDO1/2*, and *TIGIT* were also upregulated in TNBC compared to normal tissue (Figure 2D; Supplementary Figure S3B). The expression levels of the immune checkpoint gene-set were significantly higher in TNBC than in non-TNBC in both datasets (Wilcox rank-sum test, P=1.66*10^−10^, 1.45* 10^−44^ for TCGA and METABRIC, respectively). The expression levels of the immune checkpoint gene-set were also significantly higher in TNBC than in normal tissue (Wilcox rank-sum test, P=8.14*10^−11^), while showed no significant differences between non-TNBC and normal tissue (Wilcox rank-sum test, P=0.38) (Supplementary Figure S3A). Again, *TP53*-mutated TNBC had significantly higher expression levels of the immune checkpoint gene-set than *TP53*-wildtype TNBC in TCGA (Wilcox rank-sum test, P=0.04), suggesting that *TP53* mutations may have a role in the elevated expression of the immune checkpoint genes in TNBC.

In addition, Rooney *et al.* [30] identified the immunosuppressive factors that were most likely correlated with immune cytolytic activity. Strikingly, all the immunosuppressive factor genes (*C1QA*, *C1QB*, *C1QC*, *CSF2RA*, *CSF2RB*, *DOK3*, *IDO1*, *IDO2*, and *PD-L2*) were consistently upregulated in TNBC compared to non-TNBC in both datasets (Supplementary Table S8). The majority of these genes were also upregulated in TNBC compared to normal tissue including *C1QB*, *C1QC*, *DOK3*, *IDO1*, and *IDO2*.

Altogether, these results show that tumor immunosuppressive genes are likely to have higher expression levels in TNBC than in non-TNBC and normal tissue, and *TP53* mutations may result in the elevated expression of tumor immune suppressive genes in TNBC.

### TNBC has higher expression levels of many cytokine genes than non-TNBC

Cytokines are a group of small proteins that are important in the immune system [44]. Studies have shown that cytokines are important components within the TME, and play an important role in tumor immunity [45]. We compared expression levels of 261 cytokine and cytokine receptor (CCR) genes [46] between TNBC and non-TNBC (Supplementary Figure S3C, Table S9). We found that the number of CCR genes (159 in at least 1 dataset and 111 in both datasets) more highly expressed in TNBC far exceeded the number of CCR genes (42 in at least 1 dataset and 33 in both datasets) more highly expressed in non-TNBC (Fisher's exact test, P-value<2.2*10^−16^, OR=8.09). The expression levels of the CCR gene-set were significantly higher in TNBC than in non-TNBC in both datasets (Wilcox rank-sum test, P=5.5*10^−11^, 4.66*10^−43^ for TCGA and METABRIC, respectively) (Supplementary Figure S3D). The expression levels of the CCR gene-set were significantly lower in non-TNBC than in normal tissue (Wilcox rank-sum test, P=9.14*10^−12^), while showed no significant differences between TNBC and normal tissue (Wilcox rank-sum test, P=0.32) (Figure S3D).

In summary, TNBC likely has higher expression levels of CCR genes than non-TNBC. Notably, most of these cytokine receptor genes that were more highly expressed in TNBC than in non-TNBC such as *CCR1, CCR2, CCR3, CCR5, CCR7, CCR8* and *CCR9*, of which *CCR1, CCR3, CCR5, CCR7, CCR8*, and *CCR9* were also more highly expressed in TNBC than in normal tissue.

### TNBC has higher expression levels of metastasis-promoting genes than non-TNBC

In a recent study, Weyden *et al.* identified 23 genes that were involved in immune regulation of tumor metastasis [47]. Among the 23 genes, 15 (*GRSF1*, *BC017643*, *CYBB*, *FAM175B*, *BACH2*, *NCF2*, *ARHGEF1*, *FBXO7*, *TBC1D22A*, *ENTPD1*, *LRIG1*, *HSP90AA1*, *CYBA*, *NBEAL2*, and *SPNS2*) promoted tumor metastasis. Interestingly, 9 of the 15 genes showed higher expression levels in TNBC than in non-TNBC in at least 1 dataset (4 in both datasets), while only 3 genes showed higher expression levels in non-TNBC than in TNBC in at least 1 dataset (1 in both datasets) (Supplementary Figure S4, Table S10). Notably, *SPNS2* which promoted tumor metastasis *via* regulation of lymphocyte trafficking [47], had significant higher expression levels in TNBC than in non-TNBC in TCGA (expression level fold change=1.6, FDR=2.02*10^−7^) while its expression data were lacking in METABRIC. The expression levels of the metastasis-promoting gene-set were significantly higher in TNBC than in non-TNBC in both datasets (Wilcox rank-sum test, P=7.19*10^−6^, 4.82*10^−5^ for TCGA and METABRIC, respectively) (Figure 3A). TP53-mutated TNBC had significantly lower expression levels of the metastasis-promoting gene-set than *TP53*-wildtype TNBC in METABRIC (Wilcox rank-sum test, P=0.019).

**Figure 3.**
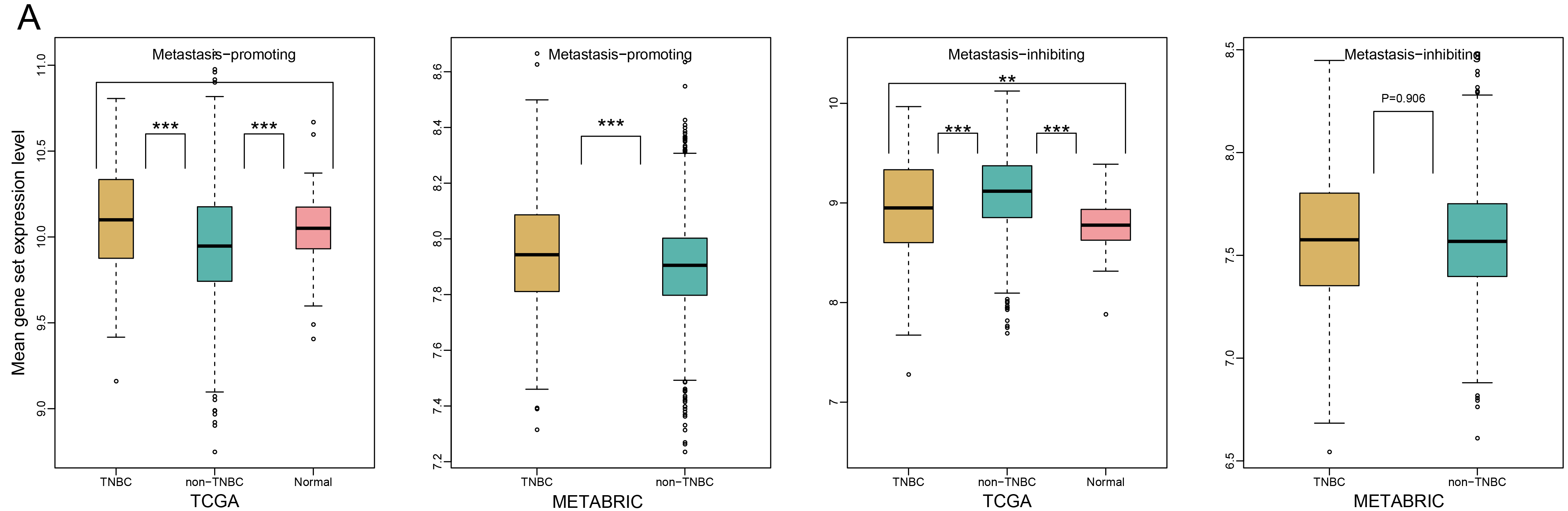
Comparison of expression levels of metastasis-promoting, metastasis-inhibiting, inflammation-promoting, and parainflammation genes between TNBC and non-TNBC. **A.** Comparison of expression levels of the metastasis-promoting and metastasis-inhibiting gene-sets between TNBC and non-TNBC. **B.** Heat-map for expression levels of inflammation-promoting genes and parainflammation genes in TNBC and non-TNBC. **C.** Comparison of expression levels of important inflammation-promoting genes between TNBC and non-TNBC in METABRIC. Red color indicates higher gene expression levels, and blue color indicates lower gene expression levels.

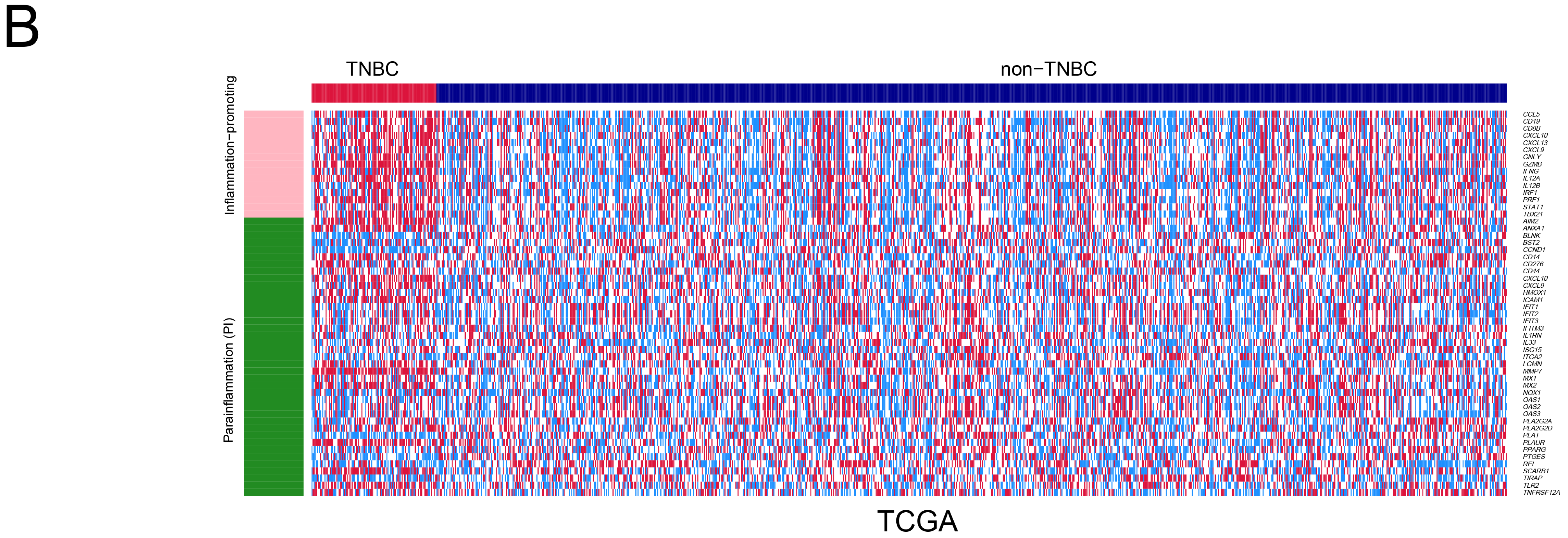

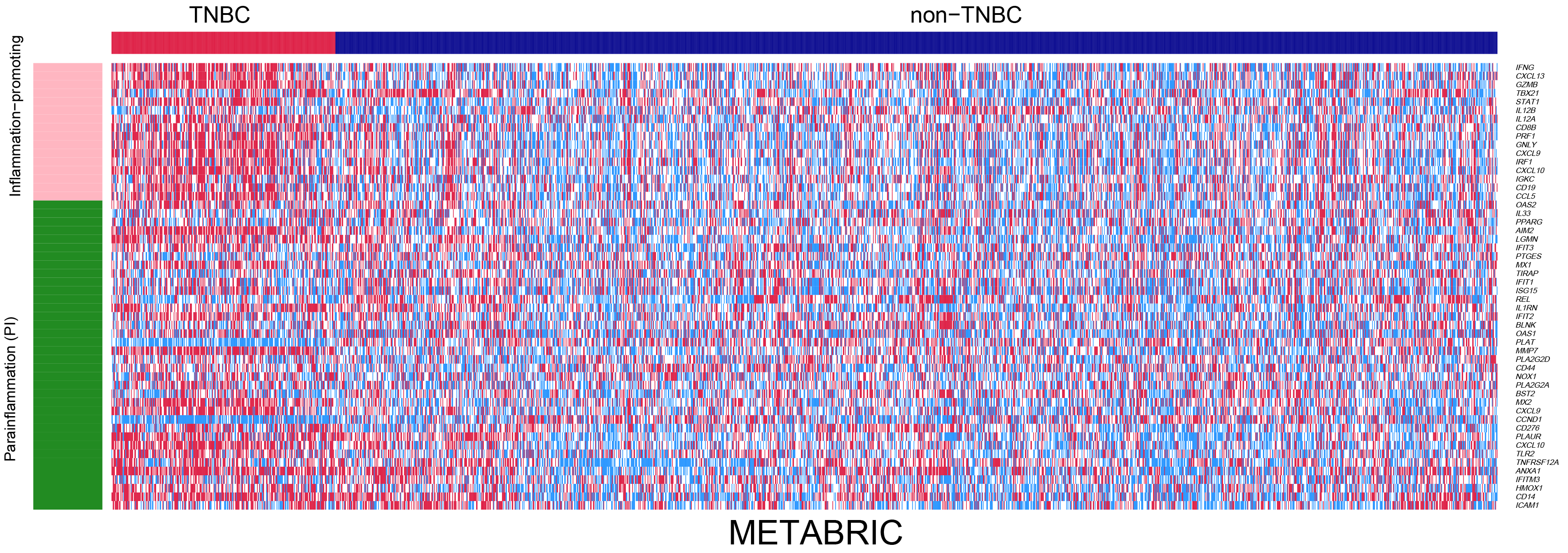

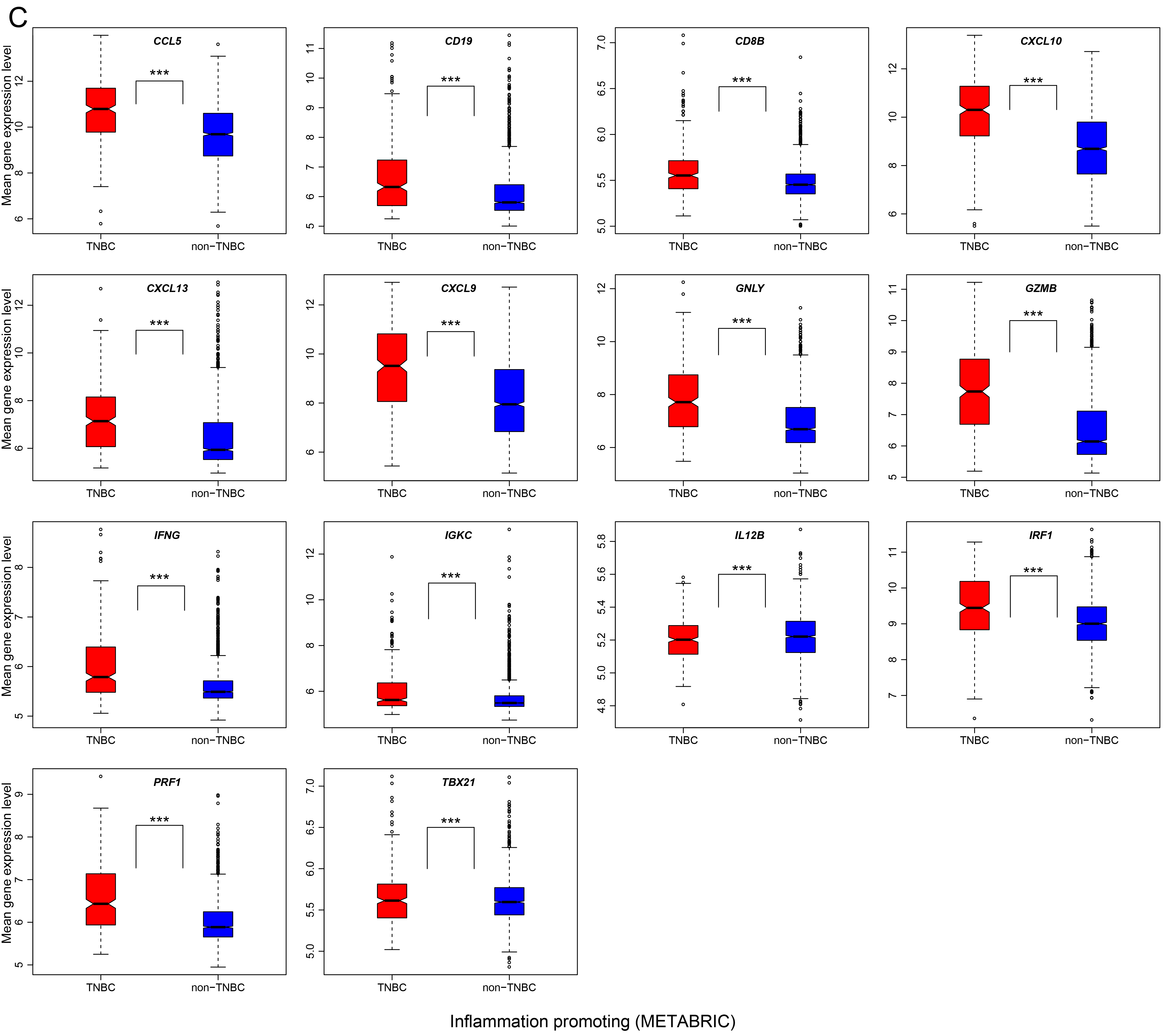

The 8 genes previously identified as inhibitors of tumor metastasis include *IRF1*, *RNF10*, *PIK3CG*, *DPH6*, *SLC9A3R2*, *IGHM*, *IRF7*, and *ABHD17A.* We compared expression levels of seven of these genes (*IGHM* had no gene expression data available in either of both datasets) between TNBC and non-TNBC, and found that 4 genes were more highly expressed in non-TNBC in at least 1 dataset (2 in both datasets), while only *IRF1* was more highly expressed in TNBC in METABRIC (Supplementary Figure S4, Table S10). The expression levels of the metastasis-inhibiting gene-set were significantly lower in TNBC than in non-TNBC in TCGA (Wilcox rank-sum test, P=2.03*10^−4^), while significantly higher in both TNBC (Wilcox rank-sum test, P=0.002) and non-TNBC (Wilcox rank-sum test, P=1.72*10^−21^) compared to normal tissue (Figure 3A).

Altogether, this data suggests that TNBC likely has elevated expression of metastasis-promoting genes and depressed expression of metastasis-inhibiting genes compared to non-TNBC, indicating that TNBC is a metastatic-prone BC subtype and this characteristic may be associated with the immune response regulation in the TME.

### TNBC has higher expression levels of inflammation-promoting genes than non-TNBC

Inflammation not only has pro-tumorigenic effects, but also influences the host immune response to tumors and cancer immunotherapy [48]. Inflammatory responses play important roles in tumor development, as seen in inflammatory BC, a rare but highly aggressive subtype of BC [49]. We compared expression levels of 16 pro-inflammatory genes [50] between TNBC and non-TNBC (Figure 3B; Supplementary Table S11). Strikingly, all 16 genes were more highly expressed in TNBC than in non-TNBC in at least 1 dataset (12 in both datasets), and 13 genes had significantly higher expression levels in TNBC than in normal tissue (Figure 3C; Supplementary Figure S5A). Notably, *STAT1* (signal transducer and activator of transcription 1) had significantly higher expression levels in TNBC compared to both non-TNBC and normal tissue. *STAT1* has been shown to play an important role in maintaining an immunosuppressive TME in BC [51]. Another gene, *GZMB* (granzyme B), together with aforementioned *GZMA*, had significantly higher expression levels in TNBC compared to both non-TNBC and normal tissue, but showed no significant expression differences between non-TNBC and normal tissue. The products of both genes are secreted by NK cells and cytotoxic T lymphocytes, and are associated with immune cytolytic activity [30]. Thus, TNBC is a BC subtype with stronger inflammatory and immune activities than the non-TNBC subtype. The expression levels of the pro-inflammatory gene-set were significantly higher in TNBC than in non-TNBC in both datasets (Wilcox rank-sum test, P=6.86*10^−12^, 6.55*10^−43^ for TCGA and METABRIC, respectively), and were significantly higher in TNBC than in normal tissue (Wilcox rank-sum test, P=4.06*10^−16^) (Supplementary Figure S5B). The expression levels of the pro-inflammatory gene-set were higher in high-grade than in low-grade TNBC (Wilcox rank-sum test, P=3.67*10^−4^), indicating that high-grade TNBC has stronger inflammatory immune response than low-grade TNBC.

Parainflammation (PI) is a low-grade inflammatory reaction that plays a role in both counteracting tumor progression and contributing to carcinogenesis [52]. In the 40 PI gene signatures [52], 27 were more highly expressed in TNBC than in non-TNBC in at least 1 dataset (14 in both datasets), compared to 10 more highly expressed in non-TNBC than in TNBC in at least 1 dataset (6 in both datasets) (Fisher's exact test, P=2.8*10^−4^, OR=6.07) (Figure 3B; Supplementary Table S11). The expression levels of the PI gene-set were significantly higher in TNBC than in non-TNBC in both datasets (Wilcox rank-sum test, P=4.81*10^−5^, 8.14*10^−31^ for TCGA and METABRIC, respectively). The expression levels of the PI gene-set were significantly higher in both TNBC and non-TNBC than in normal tissue (Wilcox rank-sum test, P=8.83*10^−8^, 0.003 for TNBC and non-TNBC, respectively) (Supplementary Figure S5B). Interestingly, *TP53*-mutated TNBC had significantly higher expression levels of the PI gene-set than *TP53*-wildtype TNBC in both datasets (Wilcox rank-sum test, P=0.04, 0.01 for TCGA and METABRIC, respectively). This confirms that PI is associated with p53 status in cancer [52]. In addition, Aran *et al.* [52] defined PI score as the single-sample gene-set enrichment analysis (ssGSEA) score [53] of the 40 PI genes for each cancer sample, and classified a TCGA cancer sample as PI positive (PI+) if the PI score was over 0.2951. Herein, TNBC had significantly higher PI scores than non-TNBC (Wilcox rank-sum test, P=0.008), and significantly higher rate of PI+ samples than non-TNBC (Fisher's exact test, P=0.02, OR=2.82). Thus, as suggested in a previous study [52], the high PI scores and high *TP53* mutation rate may cooperate to contribute to the high invasiveness of TNBC.

### TNBC with elevated expression of immune-related genes has more favorable clinical outcomes

Among the 15 gene-sets associated with immune cell types and function [30], 11 gene-sets (B cells, CD8+ T cells, macrophages, neutrophils, NK cells, pDCs, MHC class I, T cell costimulation, T cell co-inhibition, Type II IFN response, and cytolytic activity) showed significant correlation of expression levels with survival prognosis in TNBC. Strikingly, elevated expression of the 11 gene-sets was consistently associated with better OS and/or disease free survival (DFS) prognosis in TNBC (Figure 4A; Supplementary Figure S6A). Moreover, elevated expression of the HLA, Treg, immune checkpoint, immune cell infiltrate, TILs, CCR, and pro-inflammatory gene-sets was consistently associated with better OS and DFS prognosis in TNBC (Figure 4A; Supplementary Figure S6A). In addition, elevated expression of the metastasis-inhibiting gene-set was associated with better DFS prognosis in TNBC. Surprisingly, elevated expression of the metastasis-promoting gene-set was also associated with better OS and DFS prognosis in TNBC.

**Figure 4.**
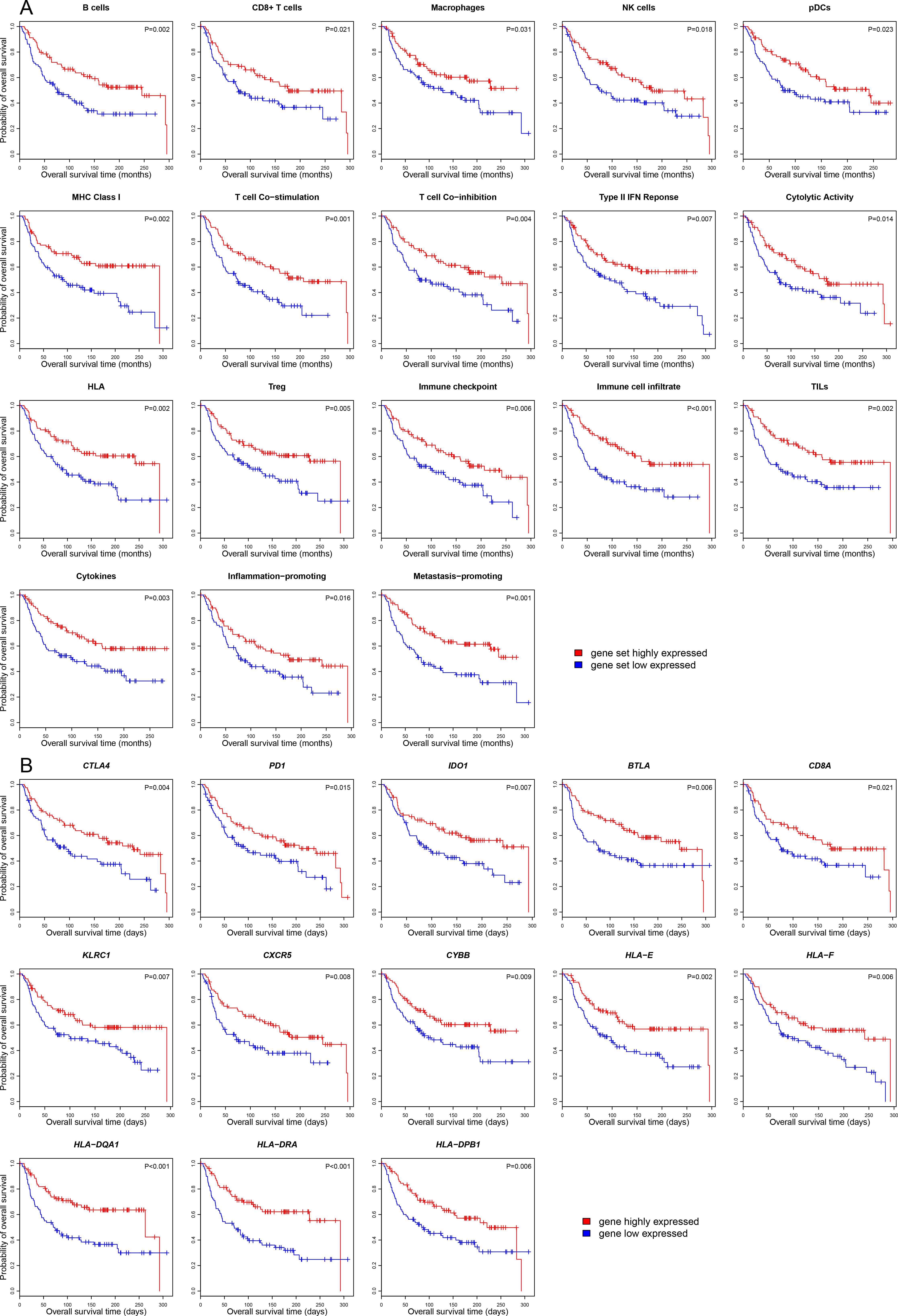
Correlation between immune gene expression and OS prognosis in TNBC. **A.** Kaplan-Meier survival curves show that elevated expression of most of the immune gene-sets is associated with better OS prognosis in TNBC (log-rank test, unadjusted *P*-value < 0.05). **B.** Kaplan-Meier survival curves show that elevated expression of a number of immune genes is associated with better OS prognosis in TNBC (log-rank test, unadjusted *P*-value < 0.05). OS: overall survival.

Interestingly, we found a substantial number of immune-related genes whose elevated expression was associated with better survival prognosis, while a few whose elevated expression was associated with worse survival prognosis in TNBC (Table 1). For example, in the 122 TILs genes, the elevated expression of 73 and 68 genes was associated with better OS and DFS prognosis in TNBC, respectively, and none was associated with worse OS or DFS prognosis in TNBC. This is consistent with prior studies showing that higher TILs densities were associated with better OS and DFS in TNBC [54, 55]. In the 47 immune checkpoint genes, the elevated expression of a number of genes was associated with better OS (17 genes) and DFS (17 genes) prognosis in TNBC, respectively. In contrast, the elevated expression of few genes was associated with worse OS (2 genes) and DFS (1 gene) prognosis in TNBC. Of the 261 CCR genes, the elevated expression of 39 genes each was associated with better OS and DFS prognosis in TNBC, respectively, compared to 5 and 3 genes whose elevated expression was associated with worse OS and DFS prognosis in TNBC, respectively. In the HLA, immune cell infiltrate, pro-inflammatory, and PI gene-sets, the elevated expression of a number of genes was associated with better OS and/or DFS prognosis, while there was no any gene whose elevated expression was associated with worse OS or DFS prognosis in TNBC. An exception was the CT gene-set in which there were 3 and 1 genes whose elevated expression was associated with better OS and DFS prognosis in TNBC, respectively, as compared to 10 and 5 genes whose elevated expression was associated with worse OS and DFS prognosis in TNBC, respectively.

We found a number of notable immune-related genes whose elevated expression was associated with better OS and DFS prognosis in TNBC such as the immune checkpoint genes *CTLA4*, *PD1*, *PD-L1*, *IDO1* and *BTLA*, cytotoxic T cell marker gene *CD8A*, NK cell marker gene *KLRC1*, Tfh cell marker gene *CXCR5*, macrophage marker gene *CYBB*, and HLA genes (Figure 4B; Supplementary Figure S6B).

**Table 1.**
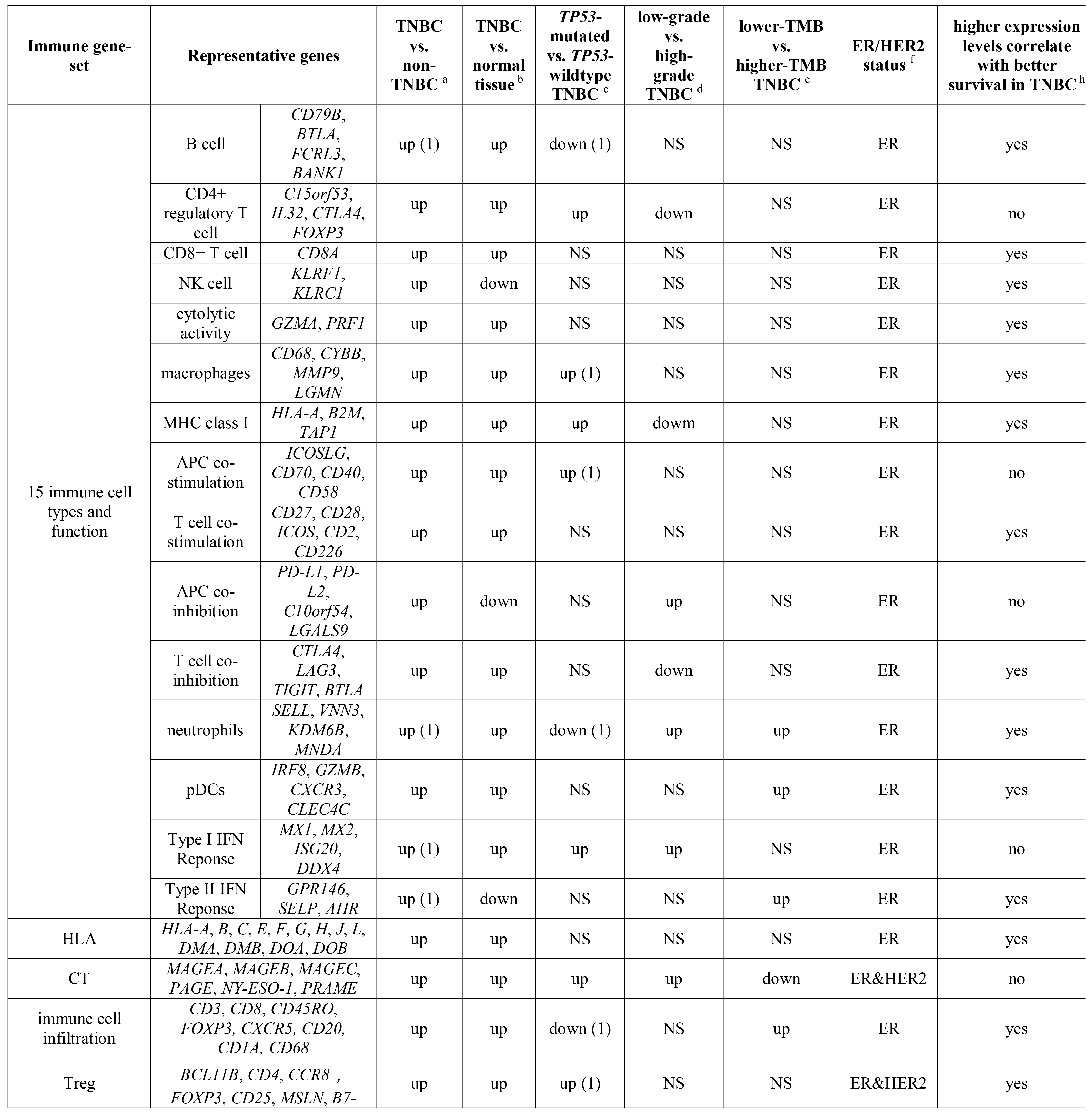
Expression of the immune-related gene-sets in TNBC.

|  |  |  |  |  |  |  |  |  |
| --- | --- | --- | --- | --- | --- | --- | --- | --- |
|  | *H4* |  |  |  |  |  |  |  |
| immune checkpoint | *CTLA4, PD1, PD-L1, PD-L2, LAG3, IDO1/2, TIGIT, BTLA* | up | up | up (1) | NS | NS | ER&ER2 | yes |
| TILs | *CD2, CD6, CD8A, CD79A, CD247, CYBB, SELL, STAT4* | up | up | NS | NS | NS | ER | yes |
| CCR | *CCR1, CCR2, CCR3, CCR5, CCR7, CCR8, CCR9, CSF2* | up | NS | NS | NS | NS | ER&HER2 | yes |
| metastasis-promoting | *SPNS2, GRSF1, BC017643, CYBB, FAM175B, BACH2, NCF2, ARHGEF1, FBXO7, TBC1D22A, ENTPD1, LRIG1, CYBA, HSP90AA1, NBEAL2* | up | NS | down (1) | NS | up | ER | yes |
| metastasis-inhibiting | *IRF1, RNF10, PIK3CG, DPH6, SLC9A3R2, IGHM, IRF7, ABHD17A* | down (1) | up | NS | NS | NS | ER or HER2<sup>g</sup> | yes (DFS) |
| pro-inflammatory | *STAT1, GZMB, CD19, CD8B, GNLY, IFNG, IL12A, PRF1* | up | up | NS | up | NS | ER | yes |
| parainflammation | *AIM2, CD14, CD276, HMOX1, LGMN, MX2, MMP7, TLR2* | up | up | up | NS | NS | ER&HER2 | no |

<sup>a</sup> The "up" indicates that the gene-set has significantly higher expression levels in TNBC than in non-TNBC in both datasets, and the "up (1)" indicates that in one of both datasets; The "down (1)" indicates that the gene-set has significantly lower expression levels in TNBC than in non-TNBC in one dataset.
<sup>b</sup> The "up" and "down" indicates that the gene-set has significantly higher and lower expression levels in TNBC than in normal tissue, respectively.
<sup>c</sup> The "up" indicates that the gene-set has significantly higher expression levels in *TP53*-mutated TNBC than in *TP53*-wildtype TNBC in both datasets, and the "up (1)" indicates that in one dataset; The "down (1)" indicates that the gene-set has significantly lower expression levels in *TP53*-mutated TNBC than in *TP53*-wildtype TNBC in one dataset.
<sup>d</sup> The "up" indicates that the gene-set has significantly higher expression levels in high-grade TNBC than in low-grade TNBC.
<sup>e</sup> The "up" and "down" indicates that the gene-set has significantly higher and lower expression levels in lower-TMB TNBC than in higher-TMB TNBC, respectively.
<sup>f</sup> Elevated expression of the immune gene-set in TNBC is associated with ER- or both ER- and HER2-status.
<sup>g</sup> The depressed expression of the metastasis-inhibiting gene-set in TNBC is associated with either ER- or HER2- status.
<sup>h</sup> The "yes" indicates that elevated expression of the gene-set is associated with better OS and DFS prognosis in TNBC, and the "yes (DFS)" indicates that elevated expression of the gene-set is associated with better DFS prognosis in TNBC.
\* The "NS" indicates no significant differences in the comparison.

### Elevated expression of immune genes in TNBC is associated with ER-or both ER-and HER2-status

The main differences in phenotypes between TNBC and non-TNBC lie in the status of ER, PR, and HER2. We explored the correlations of phenotypes with significant expression differences in the immune genes between TNBC and non-TNBC. For simplicity, we only took into account ER and HER status.

Among the 15 immune cell type and function gene-sets [30], none showed significant expression differences between TNBC and ER-/HER2+ BC in TCGA, and 4 had higher expression levels in TNBC in METABRIC. In contrast, 13 and 15 gene-sets showed higher expression levels in TNBC than in ER+/HER2-BC in TCGA and METABRIC, respectively (Supplementary Table S1). These results indicated that higher activities of the 15 immune cell types and function in TNBC were associated with the ER-status. The HLA, immune cell infiltrate, TILs, proinflammation, and metastasis-promoting gene-sets had higher expression levels in TNBC than in ER+/HER2-BC in both TCGA and METABRIC, while showed no significant expression differences between TNBC and ER-/HER2+ BC in either TCGA or METABRIC. Thus, the elevated expression of these gene-sets in TNBC was associated with the ER-status. The PI, Treg, immune checkpoint, CT, and CCR gene-sets had higher expression levels in TNBC than in ER+/HER2-BC as well as ER-/HER2+ BC in both datasets (except that PI had higher expression levels in TNBC than in ER+/HER2-BC in both datasets and than in ER-/HER2+ BC in METABRIC). Thus, the elevated expression of these gene-sets in TNBC was associated with both ER-and HER2-status. In fact, we found that almost all the immune gene-sets had significantly negative expression correlation with the ER-encoding gene *ESR1* and HER2-encoding gene *ERBB2* (Spearman correlation, FDR<0.05; Figure 5A), whereas the immune gene-sets showed stronger expression correlation with *ESR1* than with *ERBB2* (Wilcox signed-rank test, P=2.83*10^−7^, 2.98*10^−8^ for TCGA and METABRIC, respectively).

**Figure 5.**
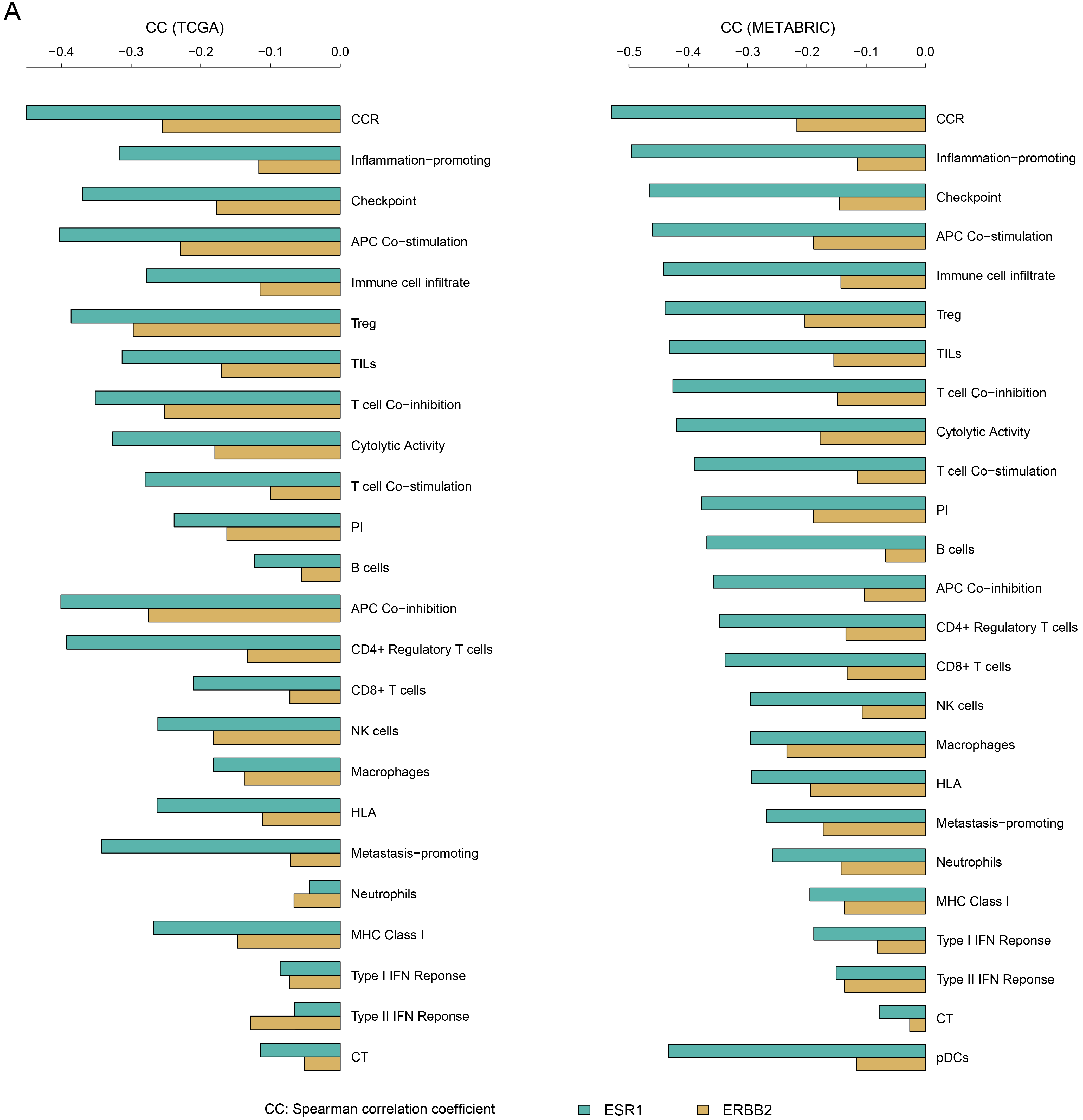
Correlation between immunogenic activity and the differential expression of genes or signaling pathways between TNBC and non-TNBC. **A.** Correlations of immunogenic activity and expression of *ESR1* and *ERBB2.* **B.** Correlations of immunogenic activity and pathway activity.

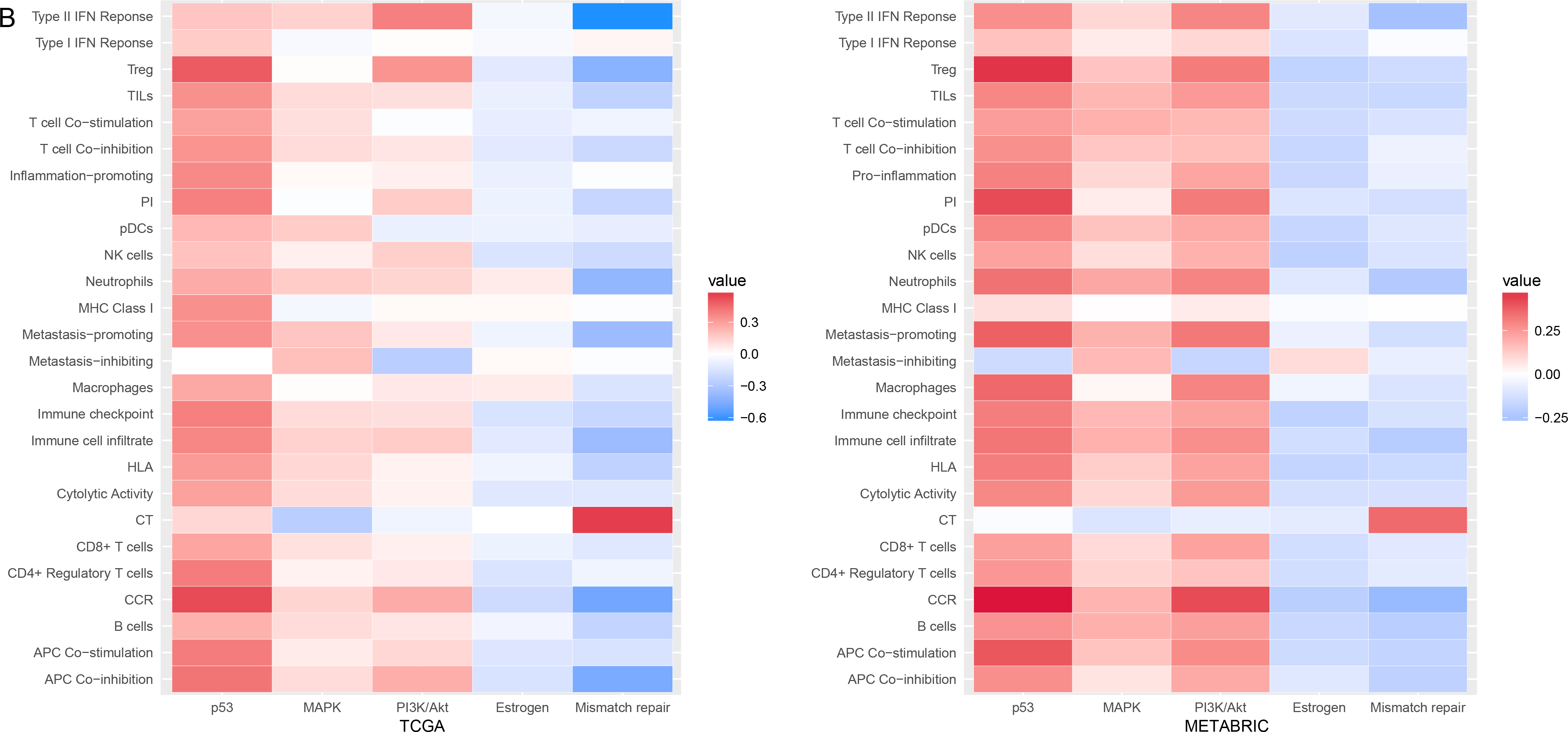

For most of the immune genes that were highly expressed in TNBC, their elevated expression was associated with the ER-status, and for some of them, their elevated expression was associated with both ER-and HER2-status (Table 1). For example, the elevated expression of the immune checkpoint genes *CTLA4*, *PD1*, *BTLA*, *TIGIT*, *VTCN1*, *CD276*, *PD-L1*, *IDO1*, and *LAG3* was associated with the ER-status since they had higher expression levels in TNBC than in ER+/HER2-BC. Among them, the elevated expression of *VTCN1* in TNBC was also associated with the HER2 status since it had higher expression levels in TNBC than in ER-/HER2+ BC in both datasets. Moreover, the elevated expression of *PD-L1*, *IDO1*, and *LAG3* in TNBC was likely associated with the HER2 status since they had higher expression levels in TNBC than in ER-/HER2+ BC in METABRIC. Furthermore, the elevated expression of a number of notable immune genes in TNBC was associated with the ER-status such as *CD4*, *CD8A*, *CSF2*, *CXCR5*, *CYBA*, *CYBB*, *GZMA*, *GZMB*, *KLRC1*, *NT5E*, *STAT1*, and *VEGFA*, and some genes whose elevated expression in TNBC was also associated with the HER2-status such as *CYBA* and *CYBB*.

In all, these results suggest that higher activities of most of the immune genes (or gene-sets) in TNBC can be attributed to the loss of ER expression, and higher activities of some immune genes (or gene-sets) can be attributed to the loss of both ER and HER2 expression.

### Distinct immunogenic activity between TNBC and non-TNBC is associated with differential signaling pathway activity

We explored the associations of immunogenic activity with the activity of 5 pathways that have significantly differential activity between TNBC and non-TNBC. The 5 pathways included the p53, MMR, PI3K/AKT, MAPK, and estrogen pathways. We selected the 5 pathways considering that *TP53* (involved in the p53 pathway) and *BRCA1* (involved in DNA MMR) had significantly higher mutation rates, and *PIK3CA* (involved in the PI3K/AKT pathway) and *MAP3K1* (involved in the MAPK pathway) had significantly lower mutation rates in TNBC than in non-TNBC concurrently in both datasets (Fisher's exact test, P<0.05). In addition, the estrogen pathway has significantly lower activity in TNBC compared to non-TNBC due to the loss of ER expression. Interestingly, we found that all the immune-related gene-sets had significantly positive correlations with the p53 pathway except the metastasis-inhibiting gene-set with a negative correlation (Figure 5B). In contrast, a majority of the immune gene-sets had significantly negative correlations with the MMR pathway, but the CT gene-set with a positive correlation (Figure 5B). As expected, almost all the immune-related gene-sets had significantly negative correlations with the estrogen pathway except the metastasis-inhibiting gene-set, which had a positive correlation (Figure 5B). In addition, a majority of the immune gene-sets had significantly positive correlations with the PI3K/AKT pathway, but the CT and metastasis-inhibiting gene-sets with a negative correlation (Figure 5B). Similarly, a majority of the immune gene-sets had significantly positive correlations with the MAPK pathway, except the CT gene-set with a negative correlation (Figure 5B). These data indicated that hyperactivation of the p53, PI3K/AKT, and MAPK pathways might promote immunogenic activity, while hyperactivation of the MMR and estrogen pathways might inhibit immunogenic activity in BC. These observations are in line with the results of previous studies [19, 56–59].

## Discussion

In the present study, we performed a comprehensive portrait of immunologic landscape of TNBC based on genomics and transcriptomics data. Strikingly, we found that all the immune-related gene-sets analyzed showed significantly higher expression levels in TNBC than in non-TNBC including 15 immune cell type and function, HLA, CT, TILs, immune cell infiltrate, Treg, immune checkpoint, CCR, metastasis-promoting, pro-inflammatory and PI gene-sets except the metastasis-inhibiting gene-set that showed significantly lower expression levels in TNBC than in non-TNBC. Our results indicated that TNBC has the strongest tumor immunogenicity of all BC subtypes. Moreover, we found that elevated expression of most of the immune-related genes (or gene-sets) in TNBC was associated with the ER-status, and that of some was associated with both ER-and HER2-status. In addition, elevated expression of the immune-related genes (or gene-sets) in TNBC was likely associated with the higher TMB in TNBC compared to non-TNBC. In fact, the higher TMB in TNBC is associated with the ER-status (Wilcox rank-sum test, P=2.32*10^−11^), but not associated with the HER2-status (Wilcox rank-sum test, P=0.29). Indeed, when we used ssGSEA score [53] instead of the gene-set mean expression levels to quantify the activity of immune cells or functions, we obtained almost the same results as those based on the gene-set mean expression level measure (Supplementary Tables S12, S13). In addition, based on the BC cell-line gene expression data from the Cancer Cell Line Project (http://www.cancerrxgene.org/), we found that 14 of the 26 gene-sets had significantly higher ssGSEA scores in TNBC cell lines than in non-TNBC cell lines. Comparatively, 4 gene-sets had higher ssGSEA scores in non-TNBC cell lines than in TNBC cell lines (Wilcox rank-sum test, P<0.05) (Supplementary Figure S7). These findings further showed that TNBC is inclined to have higher immunogenic activity than non-TNBC.

Furthermore, when we used ESTIMATE [28] to evaluate the levels of immune cell infiltration in the TME in BC, we found that TNBC had significantly higher levels of immune cell infiltration than non-TNBC in both datasets (Wilcox rank-sum test, P=3.34*10^−6^, 6.41*10^−33^ for TCGA and METABRIC, respectively) (Figure 6A). Moreover, TNBC had significantly higher levels of immune cell infiltration than ER+/HER2-BC in both datasets (Wilcox rank-sum test, P=1.14*10′^5^, 5.02*10^−39^ for TCGA and METABRIC, respectively) (Figure 6A). Compared to ER+/HER2+ BC, TNBC also had significantly higher levels of immune cell infiltration (Wilcox rank-sum test, P=5.15*10^−4^, 1.15* 10^−6^ for TCGA and METABRIC, respectively). However, TNBC showed no significantly higher levels of immune cell infiltration than ER-/HER2+ BC in either datasets (Wilcox rank-sum test, P=0.22, 0.14 for TCGA and METABRIC, respectively) (Figure 6A). This is consistent with previous studies that showed that TNBC and HER2+ BC had higher extent of immune infiltration than ER+ BC [23, 60]. In addition, we used CIBERSORT [29] to evaluate the proportions of 22 human leukocyte cell subsets within the TME in BC, and compared the proportions of each of these cell subsets between TNBC and non-TNBC. We found that activated dendritic cells, M0 macrophages, M1 macrophages, activated T cells CD4 memory, and T cells follicular helper cell subsets had significantly higher proportions in TNBC than in non-TNBC in both datasets (Wilcox rank-sum test, FDR<0.05; Figure 6B). In contrast, M2 macrophages, resting mast cells, and resting T cells CD4 memory cell subsets had significantly lower proportions in TNBC than in non-TNBC in both datasets (Wilcox rank-sum test, FDR<0.05; Figure 6B). This further demonstrates that TNBC had stronger activity of immune cells in comparison with non-TNBC. Interestingly, inflammation-inducing M1 macrophages had higher proportions in TNBC than in non-TNBC, while inflammation-inhibiting M2 macrophages that also encourage tissue repair had higher proportions in non-TNBC. This finding indicates that the TNBC disease state promotes inflammatory infiltrates and depresses tissue repair compared to non-TNBC, which may promote TNBC invasion [61, 62].

**Figure 6.**
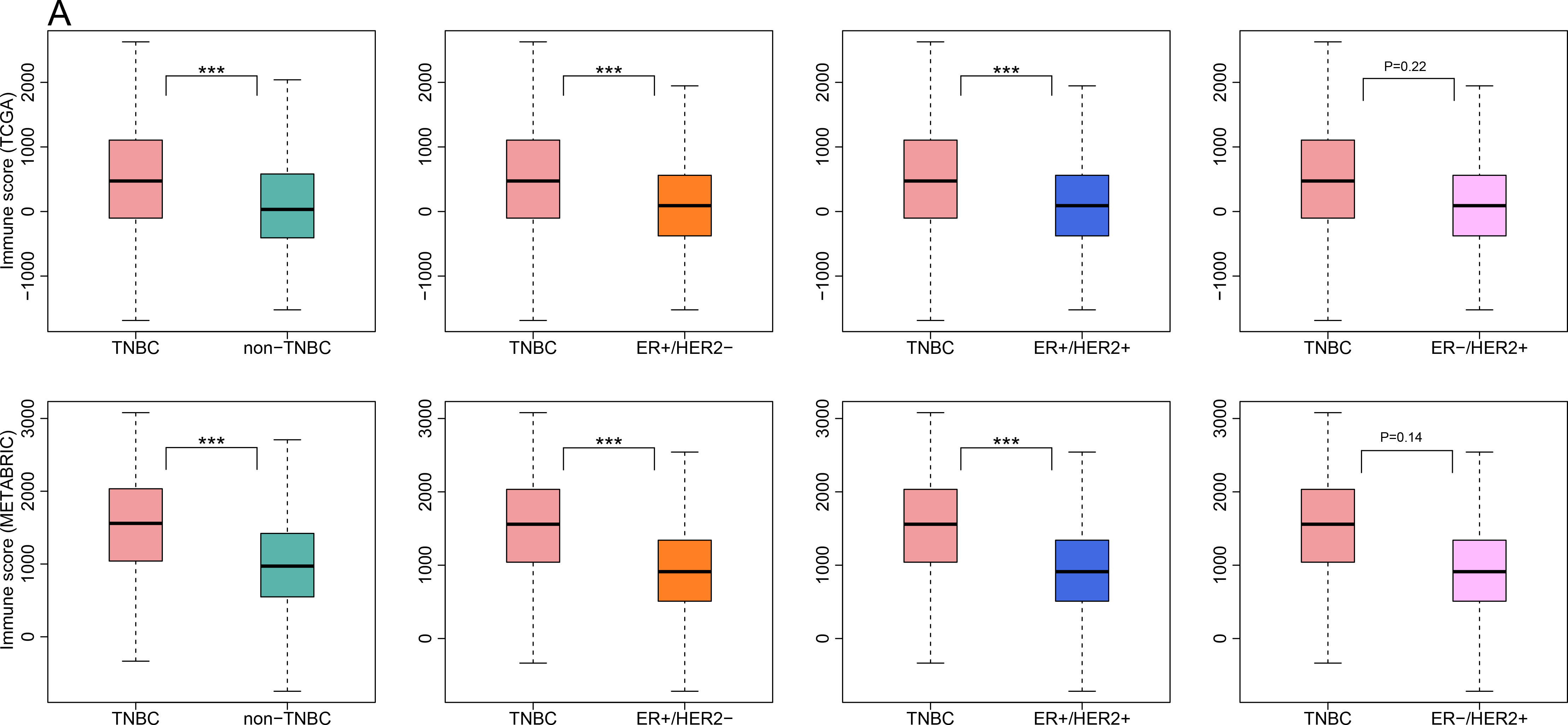
Comparison of the levels of immune cell infiltration in the tumor microenvironment between TNBC and non-TNBC. **A.** TNBC shows significantly higher degree of immune cell infiltration than non-TNBC based on ESTIMATE evaluation. **B.** TNBC has significantly different leukocyte cell subset infiltrates from non-TNBC based on CIBERSORT evaluation.

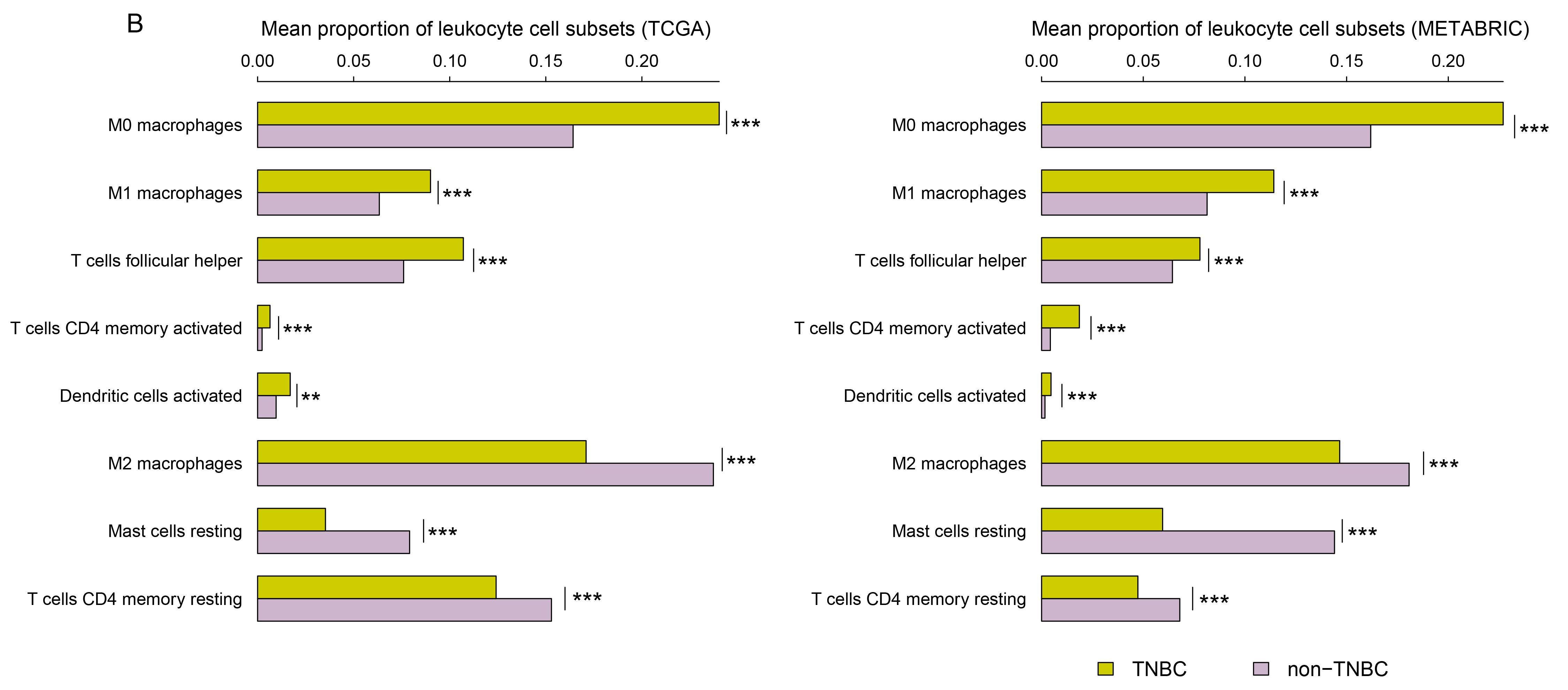

TMB has been associated with clinical response to immunotherapy [15–17]. Several cancer types with high TMB, such as melanoma [63] and NSCLC [64], have shown positive response to immune checkpoint blockade treatment. We compared expression levels of the immune gene-sets between higher-TMB and lower-TMB TNBC. We found that neutrophils, pDCs, Type II IFN response, immune cell infiltrate, and metastasis-promoting gene-sets had significantly higher expression levels in lower-TMB TNBC than in higher-TMB TNBC (Wilcox rank-sum test, P<0.05) (Supplementary Figure S8A). In contrast, the CT gene-set showed significantly higher expression levels in higher-TMB TNBC than in lower-TMB TNBC (Wilcox rank-sum test, P=0.004) (Supplementary Figure S8A). No significant expression differences between higher-TMB and lower-TMB TNBC was observed in the other gene-sets. The correlations of TMB with immune cell activities and function in TNBC should be elucidated in future studies.

Interestingly, we found that immune activities in TNBC might be associated with p53 status. Indeed, the gene-sets of CD4+ regulatory T cells, macrophages, MHC Class I, APC costimulation, Type I IFN response, CT, Treg, immune checkpoint, and PI had significantly higher expression levels in TP53-mutated TNBC than in TP53-wildtype TNBC. In contrast, B cells, neutrophils, NK cells, Type II IFN response, immune cell infiltrate, and metastasis-promoting gene-sets had significantly lower expression levels in *TP53*-mutated TNBC than in *TP53-* wildtype TNBC (Supplementary Figure S8B). These data suggest that p53 may play a role in tumor immune regulation, and p53 dysfunction may contribute to tumor immunosuppression *via* the upregulation of tumor immunosuppressive factors such as Treg and immune checkpoint genes, and downregulation of antitumor immune infiltration factors such as immune cell infiltrate genes.

Another interesting finding was that the elevated expression of most of the immune-related gene-sets was associated with better survival prognosis in TNBC. It makes sense that the elevated expression of HLA, TILs, immune cell infiltrate, CCR, and metastasis-inhibiting genes is associated with better survival prognosis in TNBC since these gene products promote anticancer immune response and inhibit tumor metastasis. Furthermore, the observation that the elevated expression of Treg, immune checkpoint, pro-inflammatory and metastasis-promoting gene-sets may be associated with better survival prognosis in TNBC may be due to the high likelihood that the elevated expression of these immunosuppressive genes can promote chemotherapy sensitivity of TNBC [60, 65]. Therefore, it is crucial to balance the gene products that inhibit tumor immunosuppression versus those that promote tumor immunopotentiation when choosing chemotherapy and immunotherapy combination in TNBC.

Interestingly, we found that TNBC had significantly higher expression levels of most of the genes targeted by immunotherapy agents in clinical use or trials or in preclinical development than non-TNBC [66]. Smyth *et al.* [66] listed 26 targets for immunotherapy agents currently used in the clinic or in clinical trials. Strikingly, 22 of the 26 target genes were more highly expressed in TNBC than in non-TNBC (Supplementary Figure S9). Moreover, 12 of the 22 genes (*TNFRSF9*, *LAG3*, *CD276*, *TNFRSF4*, *PD1*, *CTLA4*, *VEGFA*, *IDO1*, *TLR9*, *CD27*, *CSF2*, and *IL21*) were more highly expressed in TNBC than in normal tissue. Smyth *et al.* [66] also listed 12 targets for immunotherapy agents in preclinical development. Of these, 9 (*B7-H4*, *PD-1H*, *BTLA*, *CD73*, *Adenosine*, *B7-H5*, *TIGIT*, *CD96*, and *SIRPα*) were more highly expressed in TNBC than in non-TNBC. Three genes (*BTLA*, *B7-H5*, and *TIGIT*) were also more highly expressed in TNBC compared to normal tissue. These results indicate that a majority of the cancer immunotherapy agents currently used in the clinic or in clinical trials or in preclinical development may be more effective against TNBC than other BC subtypes, and may be good candidates for clinical trials for TNBC immunotherapy.

Although a number of studies have already addressed the immunogenicity of TNBC [20–24], none of these studies have performed such an exhaustive analysis of almost all types of immunogenic signatures in TNBC as in the present study. In total, we have analyzed 26 immune gene-sets including 15 immune cell type and function, HLA, CT, immune cell infiltration, Treg, immune checkpoint, TILs, CCR, metastasis-promoting, metastasis-inhibiting, pro-inflammatory, and PI gene-sets that involved 820 immune-related genes. In addition, although a number of studies have associated expression levels of immune genes with clinical outcomes in TNBC [22, 24, 36, 54, 55], none of these studies have performed a comprehensive analysis of the association between a wide variety of immunogenic signatures and clinical outcomes in TNBC as in the present study. To summarize, this study provided a solid foundation for the concept that of the various BC subtypes, TNBC likely exhibits the strongest immunogenicity.

## Conclusions

In this study, we provided a comprehensive immunologic portrait of triple-negative breast cancer based on 2 large-scale BC genomics data. Our results showed that most of the immune-related genes (or gene-sets) were more highly expressed in TNBC than in non-TNBC, suggesting that TNBC has stronger immunogenicity compared to non-TNBC. Moreover, higher expression levels of immune genes were likely correlated with better survival prognosis in TNBC. In addition, p53 status and TMB may be associated with immune activities in TNBC. These findings could have important clinical implications for TNBC immunotherapy, and warrant immunotherapeutic options for TNBC.

## Methods

### Class comparison

We normalized the TCGA BC gene expression data by base-2 log transformation, and used the original METABRIC gene expression data since they have been normalized. We compared expression levels of a single gene between two classes of samples using Student's *t* test, and compared other values between two classes of samples using the Wilcox rank-sum test. The false discovery rate (FDR) was used to adjust for multiple tests. The FDR was estimated using the Benjami and Hochberg (BH) method [67]. We used the threshold of FDR < 0.05 to identify the differentially expressed genes and gene-sets. We compared expression levels of genes or gene-sets between low-grade (Grade I-II) and high-grade (Grade III-IV) TNBC only in METABRIC, and between TNBC or non-TNBC and normal tissue only in TCGA since the other dataset had no related data available. In addition, we performed TMB and mutation counts related comparisons and analyses only in TCGA since gene somatic mutation data in TCGA were obtained by whole exome sequencing while gene somatic mutation data in METABRIC were obtained by targeted exome sequencing.

### Comparison of immune cell infiltration between TNBC and non-TNBC

We used ESTIMATE [28] to evaluate the degree of immune cell infiltration in the TME in BC. For each BC sample, we obtained an immune score to quantify the degree of immune cell infiltration in the BC tissue. We compared the immune scores between TNBC and non-TNBC using the Wilcox rank-sum test.

### Comparison of proportions of leukocyte cell subsets within the TME between TNBC and non-TNBC

We first used CIBERSORT [29] to evaluate the proportions of 22 human leukocyte cell subsets, including 7 T cell types, naïve and memory B cells, plasma cells, NK cells, and myeloid subsets. CIBERSORT was run with 1000 permutations and a threshold of P < 0.05 was the criteria for the successful deconvolution of a sample. We compared the proportions of each of the 22 leukocyte cell subsets between TNBC and non-TNBC using the Wilcox rank-sum test. We used the threshold of adjusted P-value FDR < 0.05 to identify the leukocyte cell subsets with significantly different proportions between TNBC and non-TNBC.

### Exploration of the correlation between pathways and immune gene-sets

We explored the correlation between pathways and each of the 26 immune gene-sets, respectively. We downloaded 5 gene-set collections for specific pathways (p53, MMR, estrogen, MAPK, and PI3K/AKT) from KEGG (http://www.genome.jp/kegg/). To correct for the strong correlations among these pathways, we used the first-order partial correlation to evaluate the correlations between the pathways and the immune gene-sets with the R package “ppcor” [68]. Correlations between a pathway and an immune gene-set were defined as significant if FDR was < 0.05.

### Survival analyses

We performed survival analyses of TNBC patients based on gene (or gene-set) expression data. The expression value of a gene-set was defined as the average of expression values of all the genes in the gene-set. Kaplan-Meier survival curves were used to show the survival (OS or DFS) differences between gene (or gene-set) higher-expression-level patients and lower-expression-level patients. Gene (or gene-set) higher-expression-level and lower-expression-level patients were determined by the quartile values of gene (or gene-set) expression levels. If the gene (or gene-set) expression level in a patient was higher than the third quartile value, the patient was classified as gene (or gene-set) higher-expression-level, and if was lower than the first quartile value, the patient was classified as gene (or gene-set) lower-expression-level. We used the log-rank test to calculate the significance of survival-time differences between two classes of patients with a threshold of P-value < 0.05. The survival analyses were performed only in METABRIC due to insufficient number of TNBC patients with survival data available in TCGA.

### Classification of TNBC based on TMB

For each TNBC patient, we calculated the TMB score as follows:

> *total number of truncating mutations*1.5 + total number of non-truncatingmutations*1.0*.

Truncating mutations included nonsense, frame-shift deletion, frame-shift insertion, and splice-site, while non-truncating mutations included missense, in-frame deletion, in-frame insertion, and nonstop. Silent mutations were excluded from these analyses since they do not result in an amino acid change. Truncating mutations were given a higher weight considering their higher deleterious effects on gene function than non-truncating mutations. Based on the TMB score, we classified all the TNBCs into the higher-TMB and lower-TMB classes. If the TMB score in a TNBC was higher than the median value of TMB scores, the TNBC was classified as higher-TMB; otherwise it was classified as lower-TMB.

## List of abbreviations

TNBC: Triple-Negative Breast Cancer
BC: Breast Cancer
TCGA: The Cancer Genome Atlas
BRCA: BReast invasive CarcinomA
HLA: Human Leukocyte Antigen
pDCs: plasmacytoid Dendritic Cells
iDCs: immature Dendritic Cells
MHC: Major Histocompatibility Complex
Treg: regulatory T
CT: Cancer Testis
TILs: Tumor-Infiltrating Lymphocytes
CCR: Cytokine and Cytokine Receptor
PI: Parainflammation
TMB: Tumor Mutation Burden
OS: Overall Survival
DFS: Disease Free Survival
OR: Odds Ratio
FDR: False Discovery Rate
TME: Tumor MicroEnvironment

## Declarations

### Ethics approval and consent to participate

Ethical approval was waived since we used only publicly available data and materials in this study.

### Consent for publication

Not applicable.

### Availability of data and materials

We downloaded RNA-Seq gene expression profiles (Level 3), gene somatic mutations (Level 2) and clinical data for the breast invasive carcinoma (BRCA) from the TCGA data portal (https://gdc-portal.nci.nih.gov/). The METABRIC gene expression profiles, gene somatic mutations and clinical data were downloaded from the cBioPortal website (http://www.cbioportal.org/study?id=brca_metabric#summary). For survival analyses, we used clinical data from FireBrowse (http://gdac.broadinstitute.org/) for the TCGA data, and the downloaded METABRIC clinical data. The numbers of TNBC, non-TNBC, normal tissue, ER-/HER2+, ER+/HER2-, ER+/HER2+ samples are listed in Table 2. We obtained the BC cell line gene expression profiles and clinical features data from the Cancer Cell Line Project (http://www.cancerrxgene.org/). We performed all the computational and statistical analyses using R programming (https://www.r-project.org/).

**Table 2.** Datasets used in this study.

| **Sample** | **Sample size (TCGA)** | **Sample size (Metabric)** |
| --- | --- | --- |
| TNBC | 115 | 320 |
| non-TNBC | 985 | 1660 |
| ER+/HER2- BC | 435 | 1398 |
| ER-/HER2+ BC | 41 | 139 |
| ER+/HER2+ BC | 123 | 108 |
| *TP53*-mutated TNBC | 77 | 238 |
| *TP53*-wildtype TNBC | 27 | 54 |

### Competing interests

The authors declare that they have no competing interests.

### Funding

This work was supported by startup funds awarded to XW by the China Pharmaceutical University.

### Authors' contributions

ZL performed the major data analyses, helped prepare for the manuscript and supplementary data. ML performed partial data analyses, and plotted the figures. ZJ performed partial data analyses. XW conceived the research, designed analysis strategies, performed partial data analyses, and wrote the manuscript. All the authors read and approved the final manuscript.

## Acknowledgements

Not applicable.

## Supplementary data

Table S1. Comparison of expression levels of the immune cell types and functional marker genes and gene-sets between two classes of samples

Table S2. Comparison of expression levels of the HLA genes and gene-set between two classes of samples

Table S3. Comparison of expression levels of the cancer-testis (CT) genes and gene-set between two classes of samples

Table S4. Comparison of expression levels of the tumor-infiltrating lymphocytes (TILs) genes and gene-set between two classes of samples

Table S5. Comparison of expression levels of the immune cell infiltrate genes and gene-set between two classes of samples

Table S6. Comparison of expression levels of the Treg genes and gene-set between two classes of samples

Table S7. Comparison of expression levels of the immune checkpoint genes and gene-set between two classes of samples

Table S8. Comparison of expression levels of the immunosuppressive genes and gene-set between two classes of samples

Table S9. Comparison of expression levels of the cytokine and cytokine receptor (CCR) genes and gene-set between two classes of samples

Table S10. Comparison of expression levels of the metastasis-promoting and metastasis-inhibiting genes and gene-sets between two classes of samples

Table S11. Comparison of expression levels of the inflammation-promoting and parainflammation (PI) genes and gene-sets between two classes of samples

Table S12. Comparison of expression levels (ssGSEA scores) of the immune gene-sets between two classes of samples

Table S13. Expression (ssGSEA scores) of the immune gene-sets in TNBC

**Figure S1. Comparison of expression levels of immune cell types and function, and cancer-testis genes and gene-set between TNBC and non-TNBC**. **A.** Comparison of expression levels of immune cell types and function gene-sets between TNBC and non-TNBC. *: P < 0.05; **: P < 0.01; ***: P < 0.001, and it applies to all the following box charts. **B.** Heat-map for expression levels of cancer-testis genes in TNBC and non-TNBC. **C.** Comparison of expression levels of important cancer-testis genes between TNBC and non-TNBC. **D.** Comparison of expression levels of the cancer-testis gene-set between TNBC and non-TNBC. Red color indicates higher gene expression levels, and blue color indicates lower gene expression levels.

**Figure S2. Comparison of expression levels of immune cell infiltrate genes between TNBC and non-TNBC**. **A.** Comparison of expression levels of the TILs gene-set between TNBC and non-TNBC. **B.** Comparison of expression levels of immune cell subpopulation genes between TNBC and non-TNBC in TCGA. TILs: tumor-infiltrating lymphocytes.

**Figure S3. Comparison of expression levels of the Treg, immune checkpoint, and CCR genes between TNBC and non-TNBC**. **A.** Comparison of expression levels of the Treg and immune checkpoint gene-sets between TNBC and non-TNBC. **B.** Comparison of expression levels of important immune checkpoint genes between TNBC and non-TNBC in TCGA. **C.** Heat-map for expression levels of CCR genes in TNBC and non-TNBC. CCR: cytokine and cytokine receptor. **D.** Comparison of expression levels of the CCR gene-set between TNBC and non-TNBC. Red color indicates higher gene expression levels, and blue color indicates lower gene expression levels.

**Figure S4. Heat-map for expression levels of metastasis-promoting genes and metastasis-inhibiting genes in TNBC and non-TNBC**. Red color indicates higher gene expression levels, and blue color indicates lower gene expression levels.

**Figure S5. Comparison of expression levels of the inflammation-promoting and parainflammation genes between TNBC and non-TNBC**. **A.** Comparison of expression levels of important inflammation-promoting genes and parainflammation genes between TNBC and non-TNBC in TCGA. **B.** Comparison of expression levels of the inflammation-promoting and parainflammation gene-sets between TNBC and non-TNBC.

**Figure S6. Correlation between immune gene expression and DFS prognosis in TNBC**. **A.** Kaplan-Meier survival curves show that elevated expression of most of the immune gene-sets is associated with better DFS prognosis in TNBC (log-rank test, unadjusted *P*-value < 0.05). **B.** Kaplan-Meier survival curves show that elevated expression of a number of immune genes is associated with better DFS prognosis in TNBC (log-rank test, unadjusted *P*-value < 0.05). DFS: disease free survival.

**Figure S7. 14 immune gene-sets had significantly higher ssGSEA scores in TNBC cell lines than in non-TNBC cell lines**.

**Figure S8. Associations of tumor mutation burden and** *TP53* **mutation status with immunogenic activity in TNBC**. **A.** Gene-sets that showed significant expression differences between higher-TMB and lower-TMB TNBC. **B.** Gene-sets that showed significant expression differences between *TP53*-mutated and *TP53*-wildtype TNBC (Wilcox rank-sum test, P<0.05).

**Figure S9. Heat-map for expression levels of the genes targeted by immunotherapy agents in clinical use or trials or in preclinical development in TNBC and non-TNBC**.

